# The cue-reactivity paradigm: An ensemble of networks driving attention and cognition when viewing drug and natural reward-related stimuli

**DOI:** 10.1101/2020.02.26.966549

**Authors:** Lauren D. Hill-Bowen, Michael C. Riedel, Ranjita Poudel, Taylor Salo, Jessica S. Flannery, Julia A. Camilleri, Simon B. Eickhoff, Angela R. Laird, Matthew T. Sutherland

## Abstract

**Background:** The cue-reactivity paradigm is a widely adopted neuroimaging probe engendering brain activity linked with attentional, affective, and reward processes following presentation of appetitive stimuli. Given the multiple mental operations invoked, we sought to decompose cue-related brain activity into constituent components employing emergent meta-analytic techniques when considering drug and natural reward-related cues.

**Methods:** We conducted multiple coordinate-based meta-analyses delineating *common* and *distinct* brain activity convergence across cue-reactivity studies (*N*=196 articles) involving drug (*n*=133) or natural reward-related (*n*=63) visual stimuli. Subsequently, we characterized the connectivity profiles of identified brain regions by using them as seeds in task-independent and task-dependent functional connectivity analyses. Using hierarchical clustering on these connectivity profiles, we grouped cue-related brain regions into subnetworks. Functional decoding was then employed to characterize mental operations linked with each subnetwork.

**Results:** Across all studies, *pooled* activity convergence was observed in the striatum, amygdala, thalamus, cingulate, insula, and multiple frontal, parietal, and occipital regions. *Drug-distinct* convergence (drug>natural) was observed notably in the posterior cingulate cortex (PCC), dorsolateral prefrontal cortex (dlPFC), and temporal and parietal regions, whereas *distinct natural* reward convergence (natural>drug) was observed in thalamic, insular, orbitofrontal, and occipital regions. Hierarchical clustering using each regions’ connectivity profiles identified six subnetworks, involving: 1) occipital and thalamic (lateral geniculate nucleus) regions functionally linked with early visual processing, 2) occipital-temporal regions associated with higher level visual association, 3) parietal-frontal regions linked with cognitive control mechanisms, 4) posterior and ventral insula as well as anterior cingulate cortex (ACC) functionally linked with salient event detection, 5) nucleus accumbens, PCC, precuneus, ACC, and thalamus (mediodorsal) associated with subjective valuation, and 6) bilateral amygdalae, orbitofrontal, and dorsal insula regions linked with affective processes.

**Conclusions:** These outcomes suggest multifaceted brain activity during the cue-reactivity paradigm can be decomposed into more elemental processes and indicate that while drugs of abuse usurp the brain’s natural reward processing system, some regions appear distinctly related to drug cue-reactivity (e.g., PCC, dlPFC).

## INTRODUCTION

The cue-reactivity paradigm is commonly adopted in neuroimaging research to assess neurobiological processes linked with reward, behavioral motivation, craving, and in the context of addiction, to probe the incentive salience of drug-associated stimuli [1–3]. The paradigm’s central tenet is that stimuli previously predicting receipt of drug or natural reward (e.g., food) can under certain conditions, evoke stimulus-associated responses such as urge to use drug or to eat [4]. That is, learned cues come to signal the drug or natural reward such that the cues themselves trigger arousal, anticipation, and changes in behavioral motivation. Cue-reactivity can be physiological (e.g., sweating, salivation, brain activity), symbolic-expressive (e.g., craving) [5], and/or behavioral (e.g., drug[food]-seeking, consumption) [6]. Although a relatively simple procedural design, the cue-reactivity paradigm elicits widespread brain activity across numerous regions likely involved in perceptual, attentional, memory, reward, and emotional processes. Despite this complex neurobiological response, little work has attempted to systematically and quantitatively decompose cue-related brain activity into more elemental processes or to characterize activity differences between drug-related and natural-reward cues.

Given that increased attention and responsivity to drug-related stimuli is one mechanism contributing to the development and maintenance of substance use disorders (SUDs) [7], enhanced insight into cue-reactivity’s more elemental processes has potential implications for addiction prevention and treatment. For example, alcohol dependent individuals who respond to naltrexone, a medication to reduce relapse risk, demonstrate greater pretreatment ventral striatal activity to alcohol (vs. neutral) cues, suggesting that elevated cue-reactivity may represent an endophenotype to prospectively identify those most likely to show a treatment response [8]. Even after an extended period of abstinence (e.g., years), cues are an often-cited reason for relapse [9, 10]. Relapse prevention, a cognitive-behavioral intervention, facilitates identification of high-risk situations to modify problematic behaviors that increase relapse likelihood and severity. Among these high-risk situations are those provoking craving, where learned cue associations are thought to drive drug anticipation [11, 12]. As such, continued development of second-line interventions focused on cue-induced physiological, cognitive, and behavioral reactivity holds potential to reduce recidivism rates. Systematic delineation of elemental neurobiological processes linked with cue-reactivity may provide heuristic value and expedite the evolution of second-line treatments (pharmacological or cognitive behavioral) to mitigate the impact of cues on drug-seeking and -taking behaviors.

As evidenced by prior meta-analytic work, visual drug-related (vs. control) cues evoke brain activity in visual cortices, prefrontal cortex (PFC), anterior cingulate cortex (ACC), amygdala, and the striatum among users of various substances (e.g., nicotine, alcohol, cannabis, cocaine, heroin) [12–18]. Among drug-free adults, meta-analyses focused on brain reactivity to food- and sexual-related visual stimuli have documented increased activity in the ventromedial prefrontal cortex (vmPFC), amygdala, anterior insula, mediodorsal thalamus, and striatum [19, 20]. Across these two appetitive cue domains, brain regions consistently engaged appear to be involved with visual perception processes, cognitive control (PFC), attention (ACC and insula), reward (ventral striatum [VS]), habitual learning (dorsal striatum [DS]), and emotion (amygdala). For example, increased activity and elevated extracellular dopamine concentration within mesocorticolimbic circuitry correlates with drug-seeking and -taking behaviors [21, 22], and nigrostriatal circuitry is essential for habit learning and behavioral automaticity [23]. Noteworthy, a limited window on an inverted-U shaped function of neurotransmitter (i.e., dopamine) levels within these circuits is linked with optimal reward, motivation, memory, and stress functioning, as well as effective decision-making and inhibitory control [24, 25].

The overlap of brain circuits engaged during drug and natural reward-related cue-reactivity suggests common mechanisms involved in assigning value to appetitive stimuli and transforming these subjective valuations into actions [26]. An entrenched view is that repeated drug use “hijacks” the brain’s reward system which evolved to maintain survival of the organism and species by reinforcing consummatory and procreative behaviors [22, 27, 28]. Addictive drugs produce a greater magnitude and longer-lasting concentration of synaptic dopamine than natural rewards, leading to a profound remodeling of these systems following extended use [29]. Extensive evidence supports the notion that drugs of abuse usurp natural reward mechanisms via modulation of neuronal morphology [30, 31], neurotransmitter systems [32, 33], and region-to-region functional interactions involved in reward learning [34], notably in prefrontal and striatal regions. While prior work has tended to emphasize overlapping brain regions responsive to drug and natural rewards, less often highlighted are those brain regions that may be distinctly associated with drug-cue reactivity.

Here, we expanded on prior meta-analytic work to provide enhanced insight into the *pooled, common,* and *distinct* brain regions, the putative subnetworks of regions recruited, and the elemental mental operations linked with such subnetworks during appetitive cue-reactivity. To identify convergent brain activity across and between drug and natural reward-related studies, we conducted multiple coordinate-based meta-analyses and anticipated that both cue domains would largely recruit similar brain regions. To then identify subnetworks of functionally connected brain regions, we computed both task-independent and task-dependent connectivity profiles, applied hierarchical clustering on these profiles, and expected that cue-related brain regions would form groups resembling commonly observed large-scale brain networks (e.g., visual [35], executive control [36], salience [36], subjective value [37], and default mode [38, 39] networks). Finally, to characterize the mental operations putatively linked with each cue-reactivity-related subnetwork, we employed formal behavioral decoding techniques and anticipated that such decoding would yield a collection of terms with a common interpretable cognitive theme.

## METHODS

### Cue-reactivity literature search

We conducted an iterative literature search to compile a comprehensive corpus of peer-reviewed, visual cue-reactivity, functional magnetic resonance imaging (fMRI) studies focused on drugs of abuse or natural rewards published up until August 2020. In the first iteration, we searched multiple databases, including *Google Scholar* (https://scholar.google.com) and *PubMed* (https://www.ncbi.nlm.nih.gov/pubmed), for peer-reviewed articles indexed by a combination of keywords: (“cue-reactivity” OR “drug cue” OR “natural cue”) AND (“fMRI” OR “meta-analysis” OR “GingerALE”) AND/OR (“nicotine” OR “smoking” OR “cocaine” OR “cannabis” OR “heroin” OR “alcohol” OR “sexual” OR “sex” OR “food”). In the second iteration, candidate studies were identified by reviewing the bibliographies of existing meta-analyses [3, 19, 20, 26, 40–42]. Finally, we examined the reference lists of relevant articles for potential studies not located via database searches or existing meta-analyses.

The inclusion/exclusion criteria for our meta-analyses were as follows. First, only empirical English language fMRI studies assessing drug or natural reward-related cue-reactivity using visual stimuli were included (other sensory cues [e.g., gustatory, olfactory, tactile] were not considered). Second, only studies reporting activity foci as 3D coordinates (X, Y, Z) in Talairach or Montreal Neurological Institute (MNI) stereotaxic space were included (studies involving regions of interest [ROIs] derived from a brain parcellation scheme were excluded given the absence of coordinates). Third, experiments reporting coordinates from whole-brain or small-volume corrected analyses involving a within-participant contrast of drug cues>control stimuli or natural reward-related cues>control stimuli were included. Finally, relevant information was recorded regarding participant age and sex, cue type (i.e., nicotine, alcohol, cannabis, cocaine, heroin, food, or sexual), MRI scanner field strength, and processing software (e.g., AFNI, FSL, SPM).

### Identifying cue-related brain regions: Meta-analytic procedures

To highlight convergent brain activity across and between drug and natural reward-related cue-reactivity experiments, we employed a revised version [43, 44] of the Activation Likelihood Estimation (ALE) algorithm [45, 46] as implemented in NiMARE v0.0.3, a Python package for conducting neuroimaging meta-analyses (https://nimare.readthedocs.io/en/latest/). The ALE algorithm is a voxel-wise approach for identifying statistically significant spatial convergence across a collection of study coordinates by modeling brain activity foci as 3D Gaussian probability distributions, where the distributions’ widths represent sample size variability and spatial uncertainty [43, 46, 47]. Activity foci reported by primary studies as Talairach coordinates were linearly transformed to MNI space before meta-analytic assessment [48]. The ALE algorithm first generated a set of modeled activation maps for each experimental contrast, where each voxel’s value corresponded to the maximum probability of activation. Then, the voxel-wise union of all modeled experimental contrasts was calculated yielding ALE values which quantified spatial convergence throughout the brain. These ALE values were transformed into *p*-values using a cumulative distribution function and resulting ALE maps were thresholded to highlight only voxels with *p*<0.001. Using a Monte Carlo approach, multiple comparisons correction was implemented such that a minimum cluster size threshold was determined through a set of 10,000 iterations. For each iteration, foci in the dataset were first replaced by randomly selected coordinates within a gray matter mask, ALE values were then calculated for this randomized dataset, transformed to *p*-values, thresholded at *p*<0.001, and the maximum size of supra-threshold clusters was recorded. These maximum cluster size values were used to build a null distribution. Only clusters in the original thresholded ALE map larger than the cluster size corresponding to the null distribution’s 95^th^ percentile were retained in family-wise error (FWE) corrected convergence maps reported herein. In other words, multiple comparisons corrections for all analyses were applied using a cluster-forming threshold (*p_voxel-level_* <0.001) and a cluster-extent threshold (*p_FWE-corrected_* <0.05) [49]. Surface-based and axial-slice visualization of thresholded maps were generated using NiLearn plotting tools [50].

We performed multiple ALE meta-analyses delineating common and distinct brain activity convergence when considering foci obtained from a contrast of drug or natural reward-related cue presentation relative to control stimuli presentation (i.e., drug>control, natural>control). First, to assess *<u>pooled</u>* convergence across cue domains, a meta-analysis was performed utilizing all coordinates identified across both drug and natural cue-reactivity studies thereby highlighting regions consistently showing greater activity following appetitive cue presentation relative to control stimuli (i.e., cues>control). Second, we categorized coordinates into drug (i.e., drug>control) and natural reward-related (i.e., natural>control) groupings and conducted two separate meta-analyses utilizing the same thresholding described above to elucidate convergent activity within each cue domain (**Supplemental Tables S1 & S2**). To identify regions *<u>common</u>* to both domains, a conjunction analysis was performed employing the minimum statistic [51] which identified overlapping voxels from the two thresholded ALE maps (i.e., drugs>control ⋂ natural>control). Third, we performed a meta-analytic contrast analysis to statistically compare differences in activity convergence associated with drug versus natural reward cues (i.e., drug>natural, natural>drug). To identify regions *<u>distinctly</u>* linked to each cue domain, this contrast analysis first calculated the *observed* difference in ALE statistics by subtracting the unthresholded ALE image for the drug>control contrasts from the unthresholded ALE image for natural>control contrasts. Next, we then created a null distribution of ALE difference scores to assess the statistical significance of the observed differences. To do so, we pseudo-randomly permuted the experimental contrasts between groups, calculated voxel-level difference scores, and repeated this procedure 10,000 times. For each iteration, experimental contrasts were shuffled and an equal number of contrasts to that originally observed for the drug>control and natural>control conditions were assigned to each group. Then, pseudo-ALE images were generated for these permuted groupings and subtracted. Next, voxel-level *p*-values were assigned based on a given voxel’s *observed* difference score compared to that voxel’s null distribution of pseudo-ALE difference scores (*p_FWE-corrected_* <0.05). A similar approach has been used in previous meta-analytic studies to identify statistically distinct regions within one ALE map versus another [46, 52, 53]. To exclude small regions of potential spurious differences, an additional extent-threshold of 100 contiguous voxels (arbitrarily chosen) was applied.

### Subgrouping cue-related brain regions: Functional connectivity profiles and hierarchical clustering procedures

To define cliques (i.e., subgroups) of functionally connected brain regions, we computed connectivity profiles utilizing both task-independent and task-dependent functional connectivity assessments for each ROI extracted from the *pooled* cue-reactivity meta-analysis. Given that some of the pooled cue-reactivity clusters spanned multiple anatomical regions, which may represent distinct functional nodes, we defined ROIs by generating 6-mm radius spherical seeds at the local maxima within each cluster. For this, we utilized FSL’s *cluster* command and required spherical ROIs be distanced at least 20mm from each other. ROI labels were assigned via AFNI’s *whereami* command.

#### Task-independent functional connectivity: Resting-state fMRI (rs-fMRI)

For each cue-related ROI, seed-based assessments were conducted to identify task-independent functional connectivity between the average ROI time-course and all other brain voxels. To derive robust resting-state functional connectivity (rsFC) maps, we utilized the minimally pre-processed and denoised rs-fMRI data provided by the Human Connectome Project’s (HCP; [54]) Young Adult Study (S1200 Data Release; March 1, 2017). On November 12, 2019, 150 randomly selected participants (mean±SD: 28.7±3.9 years old) were downloaded via the HCP’s Amazon Web Services Simple Storage Solution repository. The sample included 77 females (30.3±3.5 years old) and 73 males (27.1± 3.7 years). While this age difference between biological sexes was significant (*t*[149]=−5.3, *p*<0.001), it is also consistent with that noted in the full S1200 Data Release [54].

Detailed HCP data acquisition parameters can be found in consortium manuscripts [55–57] and relevant scanning parameters are briefly summarized here. Each participant underwent T1-weighted and T2-weighted structural acquisitions and four rs-fMRI acquisitions (15min each) on the 3T Siemens Connectome MRI scanner with 32-channel head coil. Structural images were collected at 0.7-mm isotropic resolution. Whole-brain EPI acquisitions were acquired with TR=720ms, TE=33.1ms, in-plane FOV=208×180mm, 72 slices, 2.0mm isotropic voxels, and multiband acceleration factor=8 [58].

The S1200 Data Release contains minimally pre-processed and denoised MRI data. The pre-processing workflow is detailed in Glasser and colleagues (2016) [59] and involved typical imaging pre-processing techniques leveraging the HCP’s high-resolution data acquisition. First, T1- and T2-weighted images were aligned, bias field corrected, and registered to MNI space. Second, the fMRI pipeline involved spatial distortions removal, volume realignment to compensate for subject motion, fMRI volume registration to structural scans, bias field reduction, normalization of functional time courses to the global average, and masking of non-brain tissue. No overt spatial smoothing was performed and care was taken by the developers to minimize inadvertent interpolation-induced smoothing. To minimize physiological and/or movement artifacts, HCP functional data were denoised using FMRIB’s ICA-based X-noiseifier (FIX; [60]) to auto-classify independent components analysis (ICA) components as either “signal” (i.e., brain activity) or “noise” (e.g., non-neuronal signals) via pattern classification of multiple spatial and temporal features. Briefly, ICA was independently performed on each functional dataset and each component’s characteristics (e.g., spatial topography, frequency band power), were evaluated by a classifier. The time-series of artifactual components were then regressed from the data, providing a “cleaned”, denoised dataset.

Using the HCP’s denoised rs-fMRI dataset, the average time course from each participant’s seed ROIs was extracted as well as the average time course across all brain voxels. Within each ROIs’ separate deconvolution (FSL’s FEAT, [61]), we entered this “global signal” time course as a regressor of no interest and spatially smoothed with a 6mm FWHM kernel. Although controversial, the global signal regressor was included given that others have demonstrated it performs better than alternative and commonly used motion-correction strategies for HCP rs-fMRI data [62]. Subject-level results for each ROI were averages across each participant’s 4, 15min. rs-fMRI runs computed using a fixed-effects analysis. A group-level, mixed-effects analysis [63] was then performed to derive rsFC maps for each ROI. Images were non-parametrically thresholded using Gaussian Random Field theory-based maximum height thresholding (voxel-level FWE-corrected at *p*<0.001) [64], which has been argued to provide more spatially specific rsFC maps when using large datasets yielding highly powered studies [65].

#### Task-dependent functional connectivity: Meta-analytic coactivation modeling (MACM)

For each cue-related ROI seed, we also conducted MACM analyses to delineate functional connectivity profiles during task-based behavioral performance. Functional connectivity within this framework reflects the coactivation of spatially distinct brain regions with the seed across numerous and varied task-based neuroimaging studies [38]. Specifically, coactivation patterns are operationalized as an above-chance convergence of activity across the neuroimaging literature demonstrating simultaneous activation with a given seed location [66]. In other words, MACM maps indicate those brain regions that are most likely to coactivate with a given ROI seed across multiple task states and behavioral domains.

To map the task-dependent coactivations for our identified cue-reactivity-related ROI seeds, we utilized Neurosynth [67], a large database of over 150,000 published stereotactic coordinates from over 14,000 fMRI studies. Neurosynth compiles (or “scrapes”) published fMRI results for all reported coordinates using an automated coordinate extraction tool. As the process is automated, fMRI studies reporting results from multiple experimental contrasts are compiled into a single coordinate set and “activation” or “deactivation” foci are not explicitly categorized. While this inherent “noise” may limit interpretations, the large amount of compiled data afforded by an automated approach (versus a more detailed, yet manually curated approach) is generally regarded to outweigh such limitations.

To generate MACM maps for each ROI, we searched the Neurosynth database for all studies reporting a coordinate within each cue-related ROI mask using NiMARE. While Neurosynth tools are available for meta-analytic assessments, we opted to use the ALE algorithm implemented within NiMARE given its documented performance regarding replication of image-based meta- and mega-analyses [68]. While the standard ALE algorithm requires participant sample sizes to generate smoothing kernels for coordinate blurring, the Neurosynth database does not capture sample size. As such, we employed a consistent 15mm FWHM kernel for all study coordinates, as this parameter value has shown the greatest correspondence between coordinate-based meta-analysis (i.e., peak activations) and “gold standard” image-based meta-analysis (i.e., whole-brain statistical images) outcomes [68]. Paralleling rsFC assessments, MACM maps were thresholded with a voxel-level FWE correction of *p*<0.001.

#### Hierarchical clustering of cue-related brain regions

We performed hierarchical clustering to subgroup identified cue-related ROIs with similar functional connectivity profiles based on their: **1)** resting-state functional connectivity patterns (i.e., task-independent), **2)** meta-analytic coactivation patterns (i.e., task-dependent), and **3)** allowing us to arrive at our final groupings, the multimodal integration of both task-independent and -dependent cross-correlation matrices. rsFC and MACM cross-correlation matrices were calculated separately using unthresholded connectivity maps to reduce the impact of sparsity associated with thresholding. Three-dimensional images representing connectivity were vectorized and concatenated to create a VxM matrix, where V was the number of voxels in the standard 2mm MNI152 brain template and M was the number of maps (i.e., ROIs). Pearson correlation coefficients were then calculated between connectivity maps for each pair-wise combination of M ROIs, yielding a new MxM correlation matrix. Then, an agglomerative hierarchical cluster tree was separately calculated for the rsFC and MACM matrices, which described how the input ROIs grouped together based on the similarity of their task-independent and -dependent maps, respectively. Hierarchical clustering assembles similar elements (i.e., cue-related ROI seeds) into clusters/cliques/subgroups in a stepwise manner, such that ROIs within a given cluster have the most similar features, yet are maximally distinct across clusters. The algorithm does so by finding two clusters that are closest together, and merging the two most similar ones until all are merged as measured by a standardized *Euclidean* ‘distance’ method and *Ward’s minimum variance* ‘linkage’ [69, 70].

Hierarchical clustering of the rsFC matrix provided cliques demonstrating similar task-independent connectivity, while clustering of the MACM correlation matrix provided cliques demonstrating similar task-dependent coactivation patterns. Given these different neuroimaging modalities, it is possible (and likely) for clustering outcomes to differ (i.e., seeds assigned to different cliques or different numbers of cliques). As such, to provide a consensus view integrating both task-independent and -dependent profiles, we again performed hierarchical clustering as described above, but now using an integrated multimodal correlation matrix combining rsFC and MACM information. The multimodal correlation matrix was calculated by averaging the respective elements across rsFC and MACM matrices. Given that the multimodal correlation matrix combined data from multiple sources, we considered the resulting cluster solution as our final groupings.

The results of the rsFC, MACM, and multimodal clustering analyses were represented visually in separate dendrograms. A dendrogram displays all variables (i.e., ROIs) entered into the clustering analysis on one axis and distance between ROIs on the other axis. Variables are joined together as clusters using branches such that the distance between two variables is indicated by the branch height on the distance axis. Similarities *between* ROIs were defined by the *Euclidean* distances between columns in each correlation matrix, representing not only one ROI’s similarity to another, but also, how similar the two ROIs’ connectivity was to all other ROIs in the cluster analysis. Subsequently, distances between clusters were defined using *Ward’s minimum variance*. *Ward’s* algorithm seeks to generate clusters by minimizing the within cluster sum of squares:

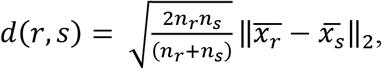

where ‖ ‖_2_ is the Euclidean distance, 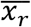 and 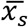 are the centroids of clusters *r* and *x*, *n_r_* and *n_s_* are the number of elements in clusters *r* and *s*. Thus, *Ward’s minimum variance values* represented similarities (or dissimilarities) between clusters which is represented on the dendrogram axes as distances.

### Functional decoding of cue-related brain region subgroups

To provide insight into the mental operations putatively linked with each cluster/clique/subgroup of cue-related ROIs from the integrated multimodal clustering solution, we performed functional decoding analyses in Neurosynth [67] leveraging the averaged and unthresholded MACM map across ROIs within a clique. Neurosynth provides distinct psychological concepts for whole-brain meta-analytic maps or identified ROIs and vice-versa. Specifically, Neurosynth computed the spatial correlation between the cliques’ input maps and maps associated with each of the 1,335 Neurosynth terms. A ranked list of maximally related psychological terms was produced providing a semi-quantitative strategy for interpreting each input map informed by the broader literature. Currently, there is no established statistical test for determining whether a term is “significantly” associated with a given input map. However, previous approaches [71, 72] have interpreted the top functional and anatomical terms, while disregarding terms which provide less interpretational value. Specifically, prior work has classified Neurosynth terms as ‘functional’, ‘anatomical’, ‘non-content’, or ‘participant-related’ (https://github.com/62442katieb/ns-v-bm-decoding). Here, we interpreted the top 10 anatomical and 10 functional terms showing the highest correlations with each cliques’ input map. Any term that designated a duplicate (or synonym) of one already identified was recorded, but not included in the final list (see **Supplemental Table S3** for details).

## RESULTS

### Literature search outcomes

We located a total of 196 peer-reviewed cue-reactivity articles, including 133 drug-related (4,093 participants; 1,243 females) and 63 natural reward-related (2,110 participants; 1,138 females) studies composed of 274 experiments/contrasts involving a total of 3,237 brain activation foci (**Supplemental Tables S1 & S2**). A PRISMA flow diagram depicting the literature search and article inclusion process is provided in **Supplemental Figure S1**. The *n*=133 drug cue-reactivity studies were further categorized by substance and included: 55 nicotine, 39 alcohol, 10 cannabis, 16 cocaine, and 13 heroin studies. Similarly, the 63-natural cue-reactivity studies were categorized by stimulus type and included: 33 sexual and 30 food studies. The total number of foci extracted from the drug and natural reward studies was 1,870 and 1,367, respectively. On average, drug cue-reactivity participants tended to be older (mean±SD: 34.4±9.1 years) and predominantly male (females: 1,243; males: 2,739 males), relative to natural reward cue-reactivity participants (28.0±7.5 years, *t*[193]=1.98, *p*<0.001), a majority of whom were female (females: 1,138; males: 972; *χ*^2^ [1, *N*=6,092]=299.0, *p*<0.001) (**Supplemental Table S1**).

### Cue-related brain regions: Meta-analytic outcomes

To first identify regions consistently demonstrating greater activity following appetitive cue presentation relative to control stimuli, we conducted a *pooled* meta-analysis utilizing foci from *both* cue-reactivity domains (i.e., cues>control). When pooling foci from both drug and natural reward-related studies, convergent brain activity was observed notably in bilateral limbic regions (amygdala, striatum, thalamus), bilateral insula, left orbitofrontal cortex (OFC), left inferior frontal, bilateral ACC, bilateral inferior occipital and parietal, left PCC, bilateral precentral, left superior parietal, and left medial occipital (**Fig. 1A; Table 1A**).

**Figure 1.**
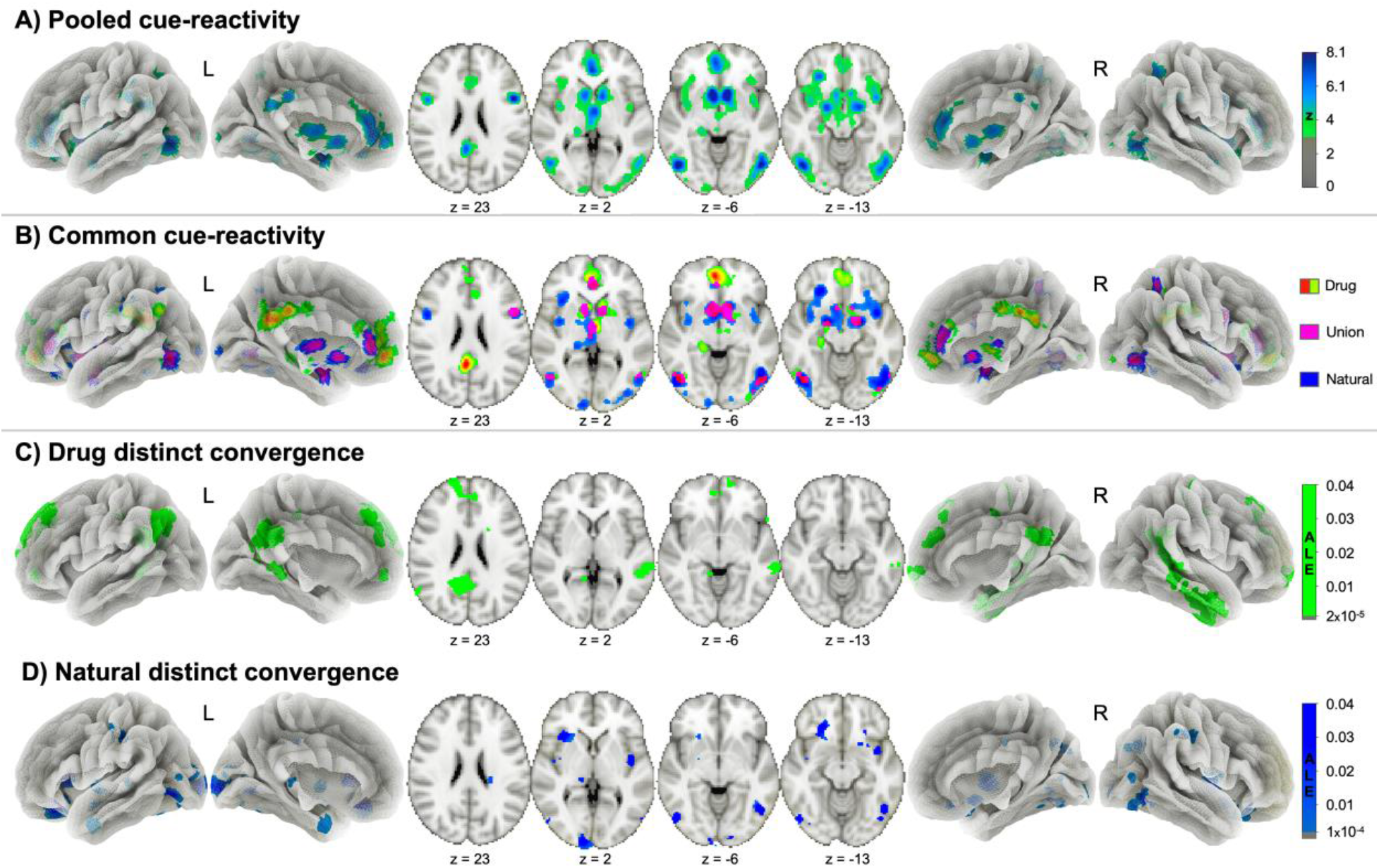
Convergent brain activity across the pooled, common, and distinct cue-reactivity meta-analytic assessments. **A)** Pooled cue-reactivity (i.e., appetitive cues > control stimuli) was observed notable in bilateral limbic (amygdala, striatum, thalamus), bilateral insula, left orbitofrontal, left inferior frontal, bilateral ACC, bilateral inferior occipital and parietal, left PCC, bilateral precentral, left superior parietal, and left medial occipital. **B)** Drug cue-reactivity (i.e., drug > control) was observed notable in left ACC, right NAc, left PCC, left amygdala, bilateral inferior occipital and parietal, and right precentral (red-green). Natural cue-reactivity (i.e., natural > control) was observed notable in left orbitofrontal, right inferior temporal and parietal, left inferior and medial occipital, right ACC, left SMG, left superior parietal, and bilateral precentral (blue). A conjunction analysis identified common areas of overlap (pink) in right caudate, bilateral ACC, left amygdala, left inferior occipital, right inferior frontal, temporal, and parietal, and left thalamus. **C)** Distinct drug cue-related convergence (i.e., drug > natural) was observed in bilateral PCC, left dlPFC, right middle and inferior temporal, left inferior parietal, and right postcentral. **D)** Distinct natural cue-related convergence (i.e., natural > drug) was observed in the bilateral orbitofrontal, multiple regions in the left occipital, right thalamus, right inferior temporal, right insula, right superior occipital, and right SMG (*p*_cluster-corrected_ <0.05, *p*_voxel_ <0.001). See **Supplemental Fig. S6** for cue-related ALE meta-analysis results using only coordinates from whole-brain assessments.

**Table 1.** Cluster coordinates for each meta-analytic assessment.

| Cluster | Region | Volume | BA | Side | X | Y | Z |
| --- | --- | --- | --- | --- | --- | --- | --- |
| *A) Pooled cue-reactivity* |  |  |  |  |  |  |  |
| 1 | amygdala | 55512 |  | L | -20 | -4 | -24 |
|  | nucleus accumbens |  |  | L | -6 | 8 | -8 |
|  | orbitofrontal gyrus |  | 11/13 | L | -26 | 32 | -16 |
|  | amygdala |  |  | R | 20 | -4 | -14 |
|  | thalamus |  |  | B | 0 | -8 | 4 |
|  | insula (ventral) |  |  | R | 38 | 8 | -12 |
|  | insula (posterior) |  |  | L | -38 | -2 | 0 |
|  | inferior frontal gyrus |  | 45 | L | -44 | 34 | 12 |
|  | insula |  |  | R | 36 | 30 | -4 |
|  | insula (dorsal) |  | 47 | L | -40 | 18 | -10 |
|  | insula (posterior) |  |  | R | 40 | -6 | 6 |
| 2 | anterior cingulate cortex | 17216 | 24 | R | 2 | 36 | 10 |
|  | anterior cingulate cortex |  | 32/33 | B | 0 | 24 | 26 |
| 3 | inferior occipital gyrus | 14992 |  | R | 48 | -70 | -8 |
|  | fusiform gyrus |  | 37 | R | 46 | -54 | -20 |
|  | inferior occipital gyrus |  |  | R | 32 | -92 | -8 |
| 4 | inferior occipital gyrus | 10512 |  | L | -46 | -68 | -12 |
|  | fusiform gyrus |  | 37 | L | -42 | -46 | -22 |
| 5 | posterior cingulate cortex | 5344 | 23 | L | -2 | -34 | 32 |
|  | precuneus |  | 7 | L | -6 | -58 | 38 |
| 6 | inferior parietal lobule | 4616 | 7 | R | 32 | -52 | 54 |
| 7 | precentral gyrus | 2944 | 6 | R | 50 | 6 | 24 |
| 8 | inferior parietal lobule | 2904 | 40 | L | -46 | -36 | 44 |
| 9 | superior parietal lobule | 2560 | 7 | L | -26 | -62 | 48 |
| 10 | precentral gyrus | 2264 | 6 | L | -46 | 4 | 28 |
| 11 | thalamus | 1760 |  | L | -20 | -32 | -2 |
| 12 | medial occipital lobe | 1512 |  | L | -12 | -98 | -2 |
| *B) Common cue-reactivity* |  |  |  |  |  |  |  |
| 1 | caudate | 7840 |  | R | 5 | 7 | -7 |
| 2 | anterior cingulate cortex | 2648 | 24 | B | 0 | 39 | 6 |
| 3 | amygdala | 2464 |  | L | -21 | -4 | -19 |
| 4 | inferior occipital gyrus | 1856 | 37 | L | -46 | -67 | -5 |
| 5 | inferior temporal gyrus | 1472 |  | R | 48 | -71 | -6 |
| 6 | inferior frontal gyrus | 1328 | 44 | R | 48 | 8 | 27 |
| 7 | thalamus | 1216 |  | L | -1 | -11 | 3 |
| 8 | inferior parietal lobule | 784 | 7 | R | 33 | -50 | 54 |
| *C) Drug distinct convergence* |  |  |  |  |  |  |  |
| 1 | dorsolateral prefrontal cortex | 9824 | 9 | L | -18 | 42 | 40 |
| 2 | posterior cingulate | 7520 | 23 | L | -4 | -50 | 26 |
| 3 | middle temporal gyrus | 5600 |  | R | 62 | -32 | -6 |
| 4 | inferior temporal gyrus | 5424 | 20 | R | 48 | -4 | -32 |
| 5 | inferior parietal lobule | 5352 |  | L | -48 | -62 | 30 |
| 6 | posterior cingulate | 1128 | 33 | R | 6 | -10 | 34 |
| 7 | postcentral gyrus | 1040 | 1/2/3 | R | 24 | -32 | 58 |
| *D) Natural distinct convergence* |  |  |  |  |  |  |  |
| 1 | orbitofrontal gyrus | 7608 | 11/13 | L | -24 | 32 | -16 |
| 2 | orbitofrontal gyrus | 1736 | 11/13 | R | 26 | 28 | -16 |
| 3 | medial occipital lobe | 1736 |  | L | -12 | -98 | -2 |
| 4 | thalamus | 1392 |  | R | 22 | -30 | 14 |
| 5 | inferior occipital gyrus | 1312 |  | L | -50 | -72 | -6 |
| 6 | inferior temporal gyrus | 1144 | 37 | R | 48 | -62 | -8 |
| 7 | middle occipital gyrus | 1120 |  | L | -26 | -92 | 8 |
| 8 | insula (posterior) | 1024 |  | R | 42 | -4 | 2 |
| 9 | superior occipital gyrus | 952 |  | R | 28 | -68 | 36 |
| 10 | supramarginal gyrus | 808 | 40 | R | 50 | -32 | 40 |

**Note.** Lettering corresponds to contrasts shown in **Figure 1: A, B, C, and D**, respectively. Cluster coordinates for A, C, and D are based on voxel peak maximum, where coordinates for B are based on center of mass. All cluster coordinates (X, Y, Z) are reported in MNI space. Volume is mm<sup>3</sup>. See **Supplemental Fig. S6** for cue-related ALE meta-analysis results using only coordinates from whole-brain assessments.

Second, we characterized convergent activity following presentation of only drug-related stimuli, only natural reward-related stimuli, and the overlapping brain regions *common* to both cue domains. The drug cue meta-analysis (i.e., drug>control) identified significant activity convergence in nine clusters, including: left ACC, right nucleus accumbens (NAc), left PCC, left amygdala, bilateral inferior occipital and parietal, and right precentral regions (**Fig. 1B**, red-green). The natural reward meta-analysis (i.e., natural>control) identified significant convergence in ten clusters, including: left OFC, right inferior temporal and parietal, left inferior and medial occipital, right ACC, left supramarginal (SMG), left superior parietal, and bilateral precentral regions (**Fig. 1B**, blue). We then performed a conjunction analysis to highlight *common* regions of convergence across both cue domains (i.e., [drug>control] AND [natural>control]) and observed overlap in eight clusters, including: right caudate, bilateral ACC, left amygdala, left inferior occipital, right inferior frontal, temporal, and parietal, and left thalamus regions (**Fig. 1B**, magenta; **Table 1B**).

Finally, to elucidate *distinct* regions of convergence (i.e., domain specificity) for drug versus natural reward-related stimuli, we performed a contrast analysis. When considering drug distinct convergence (i.e., [drug>control] > [natural>control]), we identified seven significant clusters including the bilateral PCC, left dorsolateral prefrontal cortex (dlPFC), right middle and inferior temporal, left inferior parietal, and right postcentral regions (**Fig. 1C; Table 1C**). When considering natural reward distinct convergence (i.e., [natural>control] > [drug>control]), we identified ten significant clusters, including: the bilateral OFC, multiple regions in the left occipital, right thalamus, right inferior temporal, right insula, right superior occipital, and right SMG regions (**Fig. 1D; Table 1D**).

### Subgroups of cue-related regions: Functional connectivity profiles and hierarchical clustering outcomes

To delineate task-independent and task-dependent functional connectivity profiles for cue-related ROIs identified via the *pooled* meta-analysis above, we performed seed-based rsFC and MACM assessments (**Supplemental Fig. S2 & S3**). Visual inspection of the rsFC and MACM hierarchical clustering dendrograms suggested a threshold of *n*=6 cliques for both neuroimaging modalities. Clustering solutions for task-independent rsFC patterns (**Supplemental Fig. S4**) and task-dependent MACM maps (**Supplemental Fig. S5**) yielded generally similar clique composition, with some notable differences (**Fig. 2**). Clique composition was largely consistent across both modalities when considering Clique 1 which demonstrated connectivity profiles originating in primary visual regions, including the lateral geniculate nucleus (LGN) of the thalamus, accompanied by shifts to visual association areas. Clique 2 for the rsFC solution was uniquely composed of extrastriate regions, whereas for the MACM solution, extrastriate regions clustered with the primary visual seeds in Clique 1. Clique 3 in the rsFC solution was largely consistent with Clique 2 in the MACM solution, where both included lateral parietal and frontal regions. Connectivity profiles for rsFC Cliques 4 and 5 and MACM Cliques 3, 4, and 5 were characterized by more medial and dorsal areas. Clique 4 in the rsFC solution corresponded to two distinct Cliques in the MACM solution, where MACM Clique 3 included more inferior frontal and anterior insula regions and MACM Clique 5 included more ventral and posterior insula regions. MACM Clique 4 was composed of only two regions (PCC and precuneus), whereas these regions clustered with NAc, thalamic (mediodorsal), and dorsomedial PFC regions in rsFC Clique 5. A final shift in connectivity profiles to more ventromedial regions was noted for both clustering solutions in Clique 6, where the rsFC and MACM cliques included amygdala and OFC ROIs.

**Figure 2.**
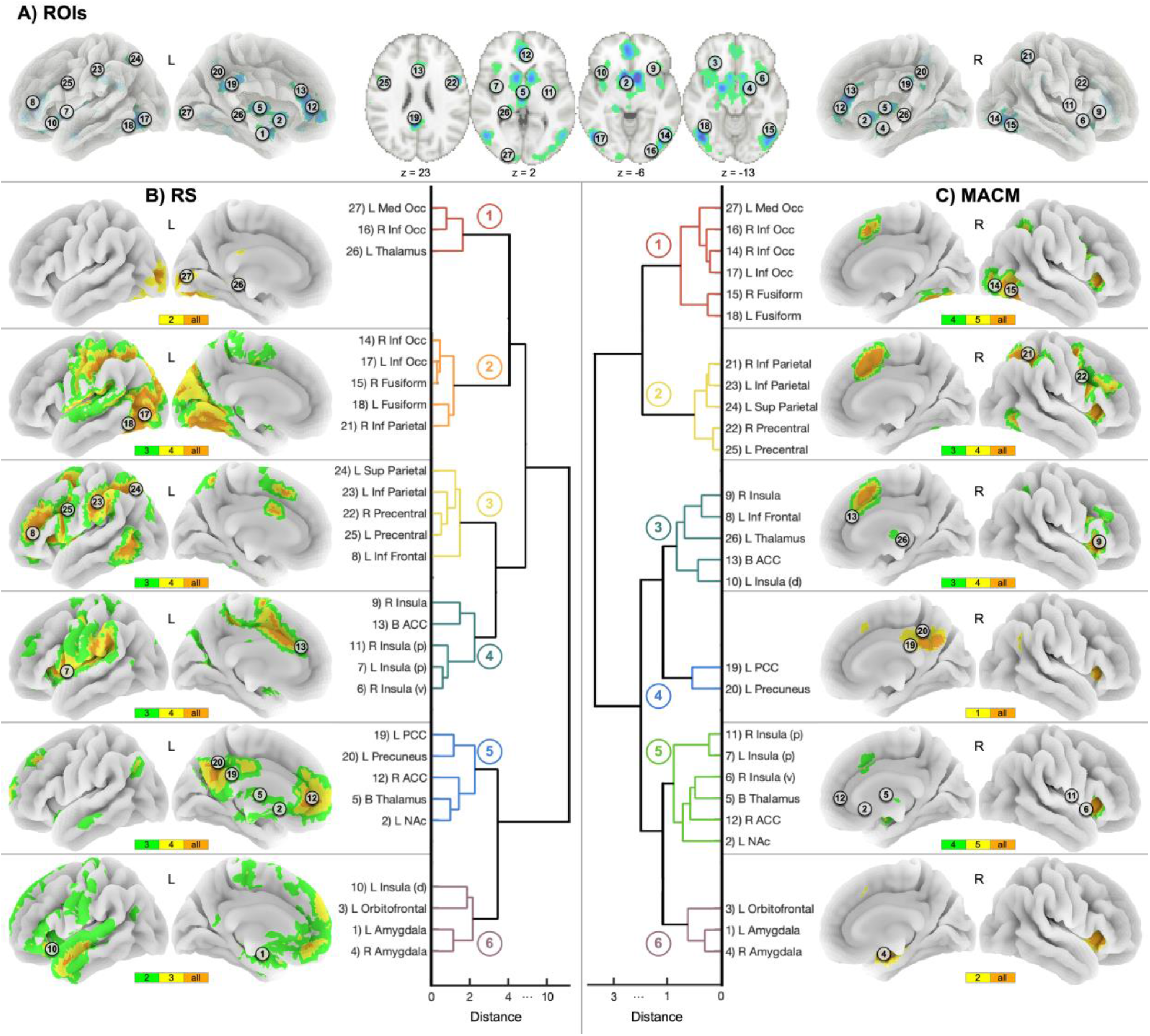
Subgroups of cue-related regions defined using task-independent (rsFC) and task-dependent (MACM) datasets. **A)** Visual representation with numberings showing cue-related ROIs. **B-C)** Hierarchical clustering of each seeds functional connectivity pattern defined subgroups of cue-related ROIs organized by functional similarity (rsFC: left, MACM: right). Both rsFC and MACM datasets yielded six cliques. The horizontal axes represent the dissimilarity (or variance) between clusters; distance was calculated using Ward’s linkage algorithm. Brain images visually represent the degree of overlap for rsFC (B) and MACM maps (C) within each clique, such that orange highlights the highest degree of overlap (all regions) and yellow and green highlights lower degrees of overlap.

To integrate both task-independent and -dependent functional connectivity profiles, we performed hierarchical clustering analysis on the integrated multimodal correlation matrix. Cue-related ROIs (**Fig. 3A**) again clustered into 6 consensus cliques (**Fig. 3B**). Clique 1 (red) consisted of left medial and right inferior occipital gyri, and left thalamus (LGN) ROIs. Clique 2 (orange) grouped together bilateral inferior occipital and bilateral fusiform gyri ROIs. Clique 3 (yellow) included bilateral inferior and left superior parietal, bilateral precentral, and left inferior frontal seeds. Clique 4 (green) consisted of multiple seeds in the bilateral insulae (posterior and ventral subregions) and the bilateral ACC. Clique 5 (blue) grouped together left PCC, left precuneus, right ACC, bilateral thalamus (mediodorsal), and left NAc ROIs. Finally, clique 6 (purple) included left insula (dorsal subregion), left OFC, and bilateral amygdalae ROIs.

**Figure 3.**
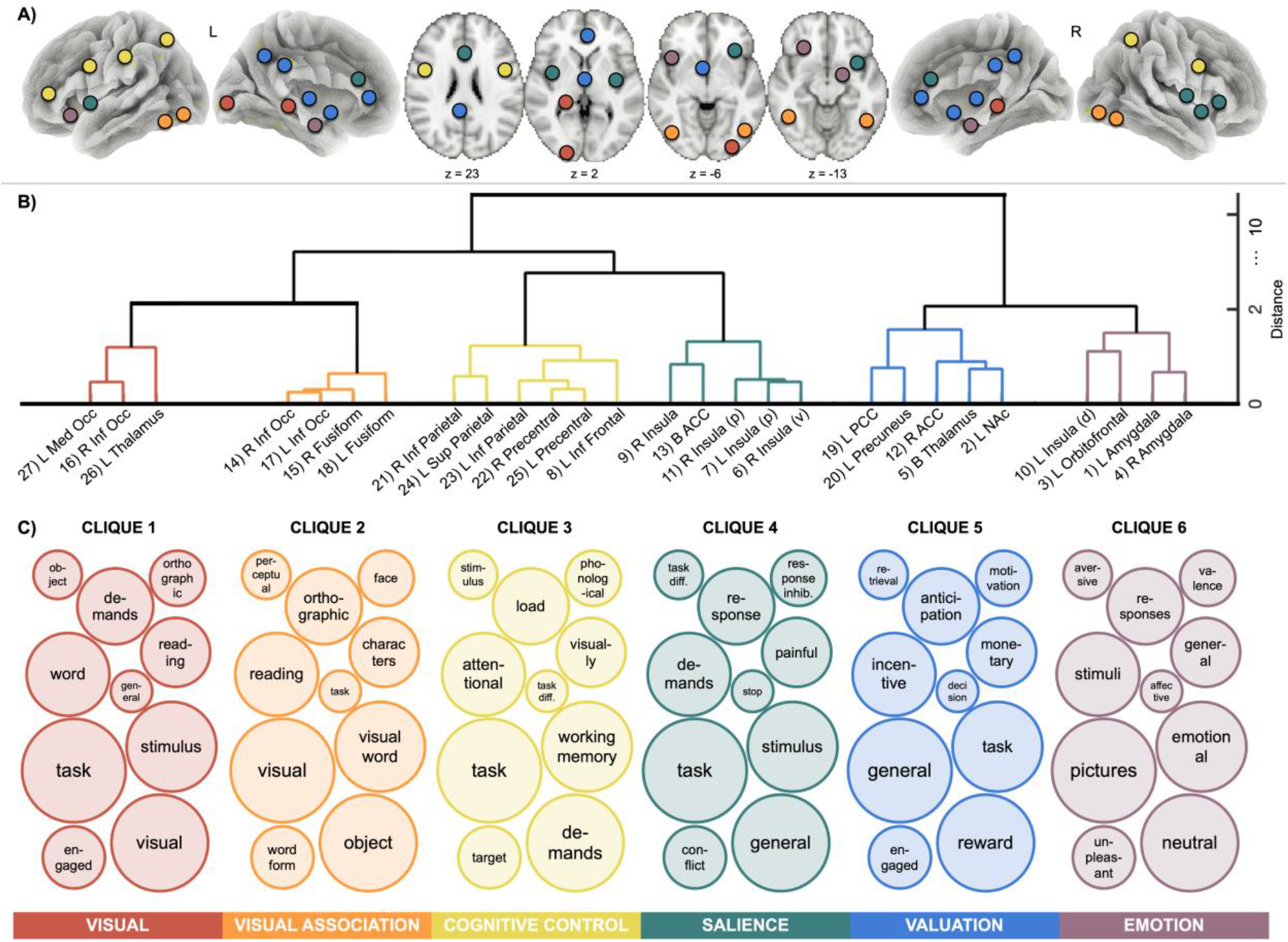
Consensus subgroups of cue-related ROIs and functional decoding linking mental operations with each clique. **A)** Visual representation of cue-related ROIs where coloring designates clique assignment. **B)** Hierarchical clustering dendrogram of the integrated multimodal clustering solution which yielded our final subgrouping of cue-related ROIs. **C)** Visual representation of each cliques’ functional decoding outcomes. Circle size designates Neurosynth’s correspondence ranking of the term with the input map, where larger circles indicate terms with higher correlation coefficients. A full list of top 10 anatomical and functional terms is located in Supplemental Information

### Functional decoding outcomes for subgroups of cue-related brain regions

We then performed functional decoding for each consensus clique to enhance insight into associated mental operations. The top 10 unique Neurosynth functional and anatomical terms with the highest correlation values for each clique were taken into consideration (**Fig. 3C**; **Supplemental Table S3**). Neurosynth decoding outcomes were used to guide the following functional interpretation of cue-reactivity-related cliques:

**Clique 1**(**Fig. 3**, **red**) was composed of occipital and thalamic (LGN) regions and was associated with the Neurosynth functional terms: *task, visual, stimulus, word, demands, reading, engaged, orthographic, object,* and *general.* These functional decoding outcomes suggested this clique was associated with the theme of <u>simple (early) visual perceptual processes</u>.
**Clique 2**(**Fig. 3**, **orange**) consisted of inferior occipital and fusiform regions and was linked with the functional terms: *visual, object, visual word, reading, orthographic, characters, word form, face, perceptual,* and *task.* These functional decoding terms suggested this clique was related to the theme of more <u>complex visual information processing and association</u>.
**Clique 3**(**Fig. 3**, **yellow**) consisted of lateral parietal and frontal regions, reminiscent of the canonical central executive network [36], and was linked with the Neurosynth terms: *task, demands, working memory, attentional, load, visually, target, phonological, stimulus*, and *task difficulty.* These outcomes suggested this clique was related to <u>cognitive control and executive functions</u>.
**Clique 4**(**Fig. 3**, **green**) was composed of ventral and posterior insulae as well as ACC regions, resembling the canonical salience network [36], and was associated with the functional terms: *task, general, stimulus, demands, response, painful, conflict, response inhibition, task difficulty*, and *stop.* These terms suggested this clique was linked with the occurrence of <u>motivationally important (i.e., salient) events</u>.
**Clique 5**(**Fig. 3**, **blue**) included ACC (ventral), PCC, thalamus, and NAc ROIs, regions often linked with the canonical default mode network [38, 39] and valuation network [37, 73, 74]. This clique was associated with the Neurosynth terms: *general, reward, task, incentive, anticipation, monetary, engaged. motivation, retrieval,* and *decision.* These outcomes suggested this clique was related to <u>subjective value and choice</u>, particularly in the context of reward.
**Clique 6**(**Fig. 3**, purple) included dorsal insula, OFC, and bilateral amygdalae regions and was linked with functional terms: *pictures, neutral, emotional, stimuli, responses, general, unpleasant, valence, aversive,* and *affective.* These functional decoding terms suggested this clique was associated with the theme of <u>emotional processing</u>.

## DISCUSSION

We decomposed cue-related brain activity into constituent components employing emergent meta-analytic techniques to provide enhanced insight into the common and distinct brain regions and, in turn, their subnetworks linked with appetitive stimuli presentation and the potential mental operations associated with such subnetworks. We first identified pooled, common, and distinct regions of convergent brain activity when considering both drug and natural reward-related cue-reactivity studies. First, when collectively considering appetitive cues, *pooled* activity convergence was observed in the striatum, amygdala, thalamus, cingulate, insula, and multiple frontal, parietal, and occipital regions. *Common* regions of convergence where drug and natural cue-reactivity overlapped included the caudate, amygdala, thalamus, ACC, and inferior frontal, parietal, temporal, and occipital regions. Drug *distinct* convergence was observed notably in the PCC, dlPFC, temporal, and parietal regions, whereas natural *distinct* convergence was observed in thalamic, insular, OFC, and occipital regions. Second, we characterized the functional connectivity profiles of those regions showing convergent activity following appetitive cue presentation by leveraging large task-independent and task-dependent MRI datasets. Lastly, we identified subnetworks/cliques of cue-reactivity-related regions from the functional connectivity profiles of ROIs and subsequently linked each subnetwork with more elemental mental operations. Based on these brain and behavioral profiles, we suggest that cue-reactivity engenders brain activity linked with visual, visual association, cognitive control, salience, valuation, and emotional operations.

### Common and distinct brain regions across cue-reactivity domains

When considering activity across both drug and natural reward-related stimuli, the *pooled* meta-analytic outcomes identified limbic (amygdala, striatum, thalamus), cingulate (ACC, PCC), insula, OFC, inferior frontal, superior parietal, precentral, and medial occipital regions of convergence across the cue-reactivity literature. Our findings largely replicate priori meta-analytic assessments regarding the recruitment of the amygdala, striatum, ACC, PCC, insula, inferior/superior parietal, precentral gyrus, and inferior/medial occipital lobe following appetitive cue presentation [26]. The primary distinction between our findings and those of Noori and colleagues’ (2016) is that they reported additional regions of convergence in inferior and middle temporal gyri. When considering *common* meta-analytic outcomes, we observed overlap between drug and natural cue-reactivity in the caudate, amygdala, thalamus, ACC, and inferior frontal, parietal, temporal, and occipital regions. Our results replicate Noori and colleagues’ (2016) findings regarding cue domain overlap in the caudate, amygdala, ACC, and inferior frontal and parietal, and extend results to the thalamus, and inferior temporal and occipital regions. These regions identified as overlapping across drug and natural reward-related cue-reactivity studies potentially represent brain regions where drugs of abuse usurp reward mechanisms evolved to maintain survival of the organism and species [75].

Moving beyond prior meta-analytic work, we identified brain regions demonstrating greater activity convergence during drug-related (vs. natural reward) cue-reactivity studies. Given we anticipated that both appetitive cue domains would largely recruit overlapping regions, we were initially surprised to observe multiple regions robustly linked with drug cue-reactivity notably including the PCC and dlPFC. Although prior meta-analytic work has not directly contrasted drug versus natural reward cue-reactivity, prior drug-specific meta-analyses have highlighted similar regions. For example, an alcohol cue-reactivity meta-analysis similarly identified PCC as well as temporal lobe convergence among studies of alcohol dependent (but not non-dependent) individuals [17]. Another previous meta-analysis focusing on cigarette cue-reactivity among daily smokers also identified activity convergence in the PCC as well as in the postcentral gyrus and superior frontal regions [14]. When considering drug-related cue reactivity irrespective of specific substances, meta-analyses have also noted activity convergence in the PCC and multiple frontal regions as well as in the amygdala, VS, inferior parietal, and occipital cortices [13]. Taken together and when combined with our direct meta-analytic contrast of drug versus natural reward-related cues, these observations highlight a potentially critical role for the PCC and dlPFC in drug-related cue-reactivity.

The PCC, a key node of the default mode network (DMN), has received relatively less attention in the neuroimaging of drug addiction field relative to, for example, the striatum or insula. The PCC is linked with internally-directed [76], or self-referential mental operations including value-based attentional capture during perceptual decision-making [77, 78], attribution of personal meaning to salient events [79], and autonomic arousal and awareness [80]. Preclinical work with macaques has demonstrated that PCC neurons signal decision outcomes in a gambling task [81]. That work also demonstrated that this reward-outcome information was maintained by PCC neurons throughout the task which ultimately predicted changes in choice behavior. Such findings suggest that PCC neurons are involved in subjective evaluative processes in reward-guided decision-making [81, 82]. This link with reward-guided decision-making may provide insight into the PCC’s critical role in addiction. For example, stroke damage to the PCC can result in the complete disruption of cigarette smoking [83]. We and others have suggested that the functional interactions between the PCC and other DMN regions may contribute to ruminations about substance use which could, in turn, perpetuate the addiction cycle [76, 84–87]. Taken together, drug-cues, more so than other appetitive cues, appear to engage the PCC which influences internally-directed processes such as value-based decision-making that may contribute to a strong motivational drive to obtain and take drugs.

In the context of addiction, the dlPFC plays a key role in top-down down-regulation of brain regions encoding the value of drug-related rewards [88], as well as itself encoding contextual information (i.e., cues, intertemporal drug availability) that may jointly modulate the experience of drug craving [89]. As such, the dlPFC has been a frequent target for transcranial direct current stimulation (tDCS) and transcranial magnetic stimulation (TMS) to reduce craving and, in turn, use of multiple substances [89, 90]. tDCS-induced dlPFC inactivation during drug cue presentation appears to attenuate craving via modulation of other regions including the medial orbitofrontal cortex (mOFC), ACC, and VS [89]. Whereas the mOFC is believed to track the subjective value of drug stimuli, the dlPFC is thought to integrate information regarding cue information and temporal availability which can modulate the mOFC’s value signal [91]. Our results provide further corroborating support for the critical role of the dlPFC during drug cue exposure and, in turn, drug-seeking and -taking.

### Distributed networks of cue-reactivity

SUDs are often conceptualized as impacting multiple brain circuits and networks, rather than a specific “lesion” within circumscribed brain regions [92]. Brain networks are characterized by a collection of regions, where the dynamic interactions of nodes within a network, as well as between other large-scale networks are linked with various mental operations such as: self-referential cognition (DMN), detection of and orientation to salient external and internal events [salience network (SN)], and higher-level cognition including working memory and attentional control [central executive network (CEN)] [93, 94]. As such, we examined the functional connectivity profiles (i.e., both rsFC and MACM) of cue-reactivity-related regions to cluster them into subnetworks and to delineate the more elemental mental operations associated with each subnetwork. We demonstrated that specific subnetworks within the broad collection of regions responsive to appetitive cues appear to show functional specialization. Through data-driven techniques, we identified six cliques with functional decoding outcomes indicating roles in visual, visual association, cognitive control, salience, valuation, and emotional processing.

### Visual perception subnetworks (Cliques 1 and 2)

Inherent to the cue-reactivity paradigm is the recruitment of primary (clique 1) and associative visual cortices (clique 2) linked with stimulus perception. Noteworthy, we observed activity convergence in brain regions linked with visual perceptual processes when considering appetitive (vs. control) cues, suggesting that learned associations between predictive stimuli and rewards modulate top-down attention. As evidenced by prior work, the representation of basic visual features (e.g., local contrast, location, spatial frequency) can be modulated by top-down attention and learned associations between stimuli and reward [15, 104]. Pairing visual stimuli with a reward improves stimulus detection [105, 106], reduces response times [107, 108], and increases accuracy [109]. The neural mechanism by which rewards may regulate plasticity of the visual representation of reward-predicting stimuli is through dopamine signaling [110]. Accordingly, given learned associations, chronic substance users may demonstrate increased activity within visual processing circuitry when presented with drug-related stimuli which could be conceptualized as a neurobiological manifestation of attentional bias. While sensory processes have historically been overlooked in the addiction literature, these systems may have important implications for the development and/or maintenance of substance use. Indeed, cue-elicited activation of visual processing regions is linked with clinically-relevant outcomes such as drug craving [111], craving resistance [112], dependence severity and automatic motor responses to cues [113, 114], self-recognition of substance use problems [115], and importantly, to relapse itself [116].

### Tripartite network integration (Cliques 3, 4, 5)

The tripartite network heuristic framework focuses on three large-scale brain networks [93], the DMN, SN, and CEN. The CEN, a frontoparietal system with primary nodes in the dlPFC and lateral posterior parietal cortices, is linked with exogenous, attentionally driven cognitive functions [95, 96]. The DMN, centered on nodes in the PCC, mPFC, medial temporal lobe, and angular gyrus, is typically deactivated during stimulus-driven cognitive tasks and, on the other hand, is implicated in ruminations, mind wandering, and reflections on the past [77, 97]. The SN, anchored in the dorsal ACC and frontoinsular cortices, is typically associated with orienting attention to internal or external stimuli [36, 98]. Cue-reactivity Cliques 3, 4, and 5 identified in the current study closely corresponded to these canonical large-scale networks.

Dysregulated activity within and between these three networks has been highlighted across various neuropsychiatric disorders including addiction [84, 94]. As the brain is continuously inundated with information arising from external and internal stimuli, optimal behavior necessitates control mechanisms to orient, identify, and act upon the currently most salient stimuli. The SN is often regarded as serving this role by influencing moment-to-moment information processing and “toggling” between the internally directed DMN and the externally directed CEN [84]. Across stages of the addiction cycle (i.e., taking, withdrawal, urge), the insula is often highlighted as a critical region, that through its interactions with other brain regions alters affective states (e.g., irritability), motivation (e.g., cue-reactivity), and attention (e.g., bias to drug-related stimuli) [99]. For example, during withdrawal, increased insula engagement and coupling with the DMN may serve to orient attention toward this internal physiological state at the expense of externally focused mental operations linked with the CEN [84, 86, 95]. This anticorrelation between DMN and CEN dynamics is relevant for optimal task performance, where increased DMN activation and reduced CEN activation is often linked with suboptimal task performance among healthy individuals and those diagnosed with various neuropsychiatric conditions (including SUDs) [93, 100, 101]. One of the more frequently highlighted functions of the DMN in the context of neuropsychiatric conditions is ruminations about one’s internal state and self [102, 103]. Aberrant connectivity within the DMN is linked with impaired self-awareness, negative emotions, and obsessive thoughts about drugs which may contribute to relapse and compulsive drug-taking despite negative consequences [76]. Our results provide evidence that engagement of these three large-scale networks, comprising the tripartite network, are engaged during appetitive cue-reactivity and potentially linked with self-referential cognition (DMN), detection and orientation to salient events (SN), and higher-level cognition (CEN).

### Emotional processing (Clique 6)

The final clique we identified as an elemental component of cue-reactivity grouped together amygdala, insula, and OFC regions. The amygdala can be subdivided into at least two distinct components, each comprising multiple nuclei, with cooperating functions: the basolateral complex (BLA) that encodes sensory-specific features of emotional events (i.e., Pavlovian conditioning), and the central nucleus (CeN) that encodes more general affective and motivationally significant aspects of emotional events [117, 118]. The coordination of these two components supports the amygdala’s role in both appetitive and aversive motivational systems [119]. The reciprocal connections between the BLA and the OFC and insula, are critical for regulating goal-directed behaviors [120] and modulating decision-making processes [121]. For example, in the context of reward-learning, BLA and OFC interactions are critical for behavioral responses dependent on the acquisition and use of conditioned stimulus-reward associations [121], with BLA and insula connections facilitating encoding and retrieval of sensory information (i.e., gustation and taste) in relation to bodily state, thereby shaping perceived valence [122]. Dysregulated affective neurocircuitry is prominently featured in theorizing regarding the development and maintenance of substance use [123, 124]. For example, in chronic drug-dependence, the amygdala is linked with aversive emotional states underlying withdrawal that through negative reinforcement, partly motivates the compulsive seeking and taking of drugs [125]. In other words, while the amygdala and OFC play prominent roles in drug-related aversive behaviors, this circuitry is also engaged when encountering drug-associated stimuli (i.e., people, places, and objects) [126]. Thus affective-related neurocircuitry potentially contributes to both positive and negative reinforcement mechanisms perpetuating drug use.

### Limitations

Potential limitations warrant attention. First, our meta-analysis was limited to functional neuroimaging experiments, thus precluding any interpretation of underlying synaptic or molecular mechanisms. Second, all meta-analyses are susceptible to biases across the literature, are limited by the primary studies’ designs, and only include significant peak activations reported by those primary studies (i.e., publication bias). Third, the ALE meta-analytic algorithm does not take into consideration the size of the cluster identified from the primary studies, resulting in less precise representations than image-based meta-analytic approaches [68]. Fourth, when utilizing data-driven analytic approaches multiple parameter values often must be specified [127], and the hierarchical clustering employed herein involved selection of two such parameters, a distance measure and a linkage algorithm. Based on prior work [69, 70, 128], we selected the standardized *Euclidean* ‘distance’ method and *Ward’s minimum variance* ‘linkage’, how the outcomes reported herein would differ as a function of the distance and linkage methods employed was beyond the scope of this work. Finally, a biological sex discrepancy among participants between appetitive cue domains was noted, where drug-related articles assessed about twice as many male relative to female participants; for the natural-reward studies biological sex was more evenly balanced. This discrepancy potentially reflects the notion that males are more likely to use most types of illicit drugs [129] and typically present with higher rates of use and dependence [130]. Differences in brain reactivity to appetitive cues as a function of biological sex remains an important area of future investigation.

### Conclusions

In sum, the current study employed emergent neuroimaging meta-analytic techniques to enhance insight into the brain regions, subnetworks of regions recruited, and more precise elemental mental operations linked with such subnetworks during appetitive cue-reactivity. We identified convergent brain activity across and between drug and natural reward-related studies via multiple coordinate-based meta-analyses. Our outcomes indicated that while drugs of abuse to some degree usurp the brain’s natural reward processing system, some regions appear distinctly related to drug cue-reactivity (e.g., PCC, dlFPC). We also identified subnetworks of functionally connected brain regions by leveraging large task-independent and task-dependent MRI datasets, and applying hierarchical clustering on these connectivity profiles. We linked each subnetwork with more elemental mental operations and suggest that cue-reactivity engenders brain activity linked with visual processing networks, the tripartite network model, and an emotion related network. Enhanced insight into the more elemental neurobiological processes engendered during appetitive cue-reactivity may provide heuristic value in the service of advancing the evolution of second-line cognitive behavioral and/or pharmacological interventions to reduce cue-induced relapse.

## ACKNOWLEDGEMENTS

Primary funding for this project was provided by NIH R01 DA041353; additional support was provided by NSF 1631325, NIH U01 DA041156, NSF CNS 1532061, NIH K01 DA037819, NIH U54 MD012393. Additional funding was provided by the Deutsche Forschungsgemeinschaft (DFG, EI 816/11-1), NIH R01-MH074457, the Helmholtz Portfolio Theme “Supercomputing and Modeling for the Human Brain” and the European Union’s Horizon 2020 Research and Innovation Programme under Grant Agreement No. 720270 (HBP SGA1) 785907 (HBP SGA2).

