## Supplemental Figures for "The cue-reactivity paradigm: An ensemble of networks driving attention and cognition when viewing drug and natural reward-related stimuli"

**SUPPLEMENTAL CONTENT**

SUPPLEMENTAL FIGURES

**Figure S1.** PRISMA flow diagram

**Figure S2.** rsFC maps for identified cue-related regions

**Figure S3.** MACM maps for identified cue-related regions

**Figure S4.** rsFC hierarchical clustering solution

**Figure S5.** MACM hierarchical clustering solution

**Figure S6.** Cue-related ALE meta-analysis results using only coordinates from whole-brain assessments

SUPPLEMENTAL REFERENCES

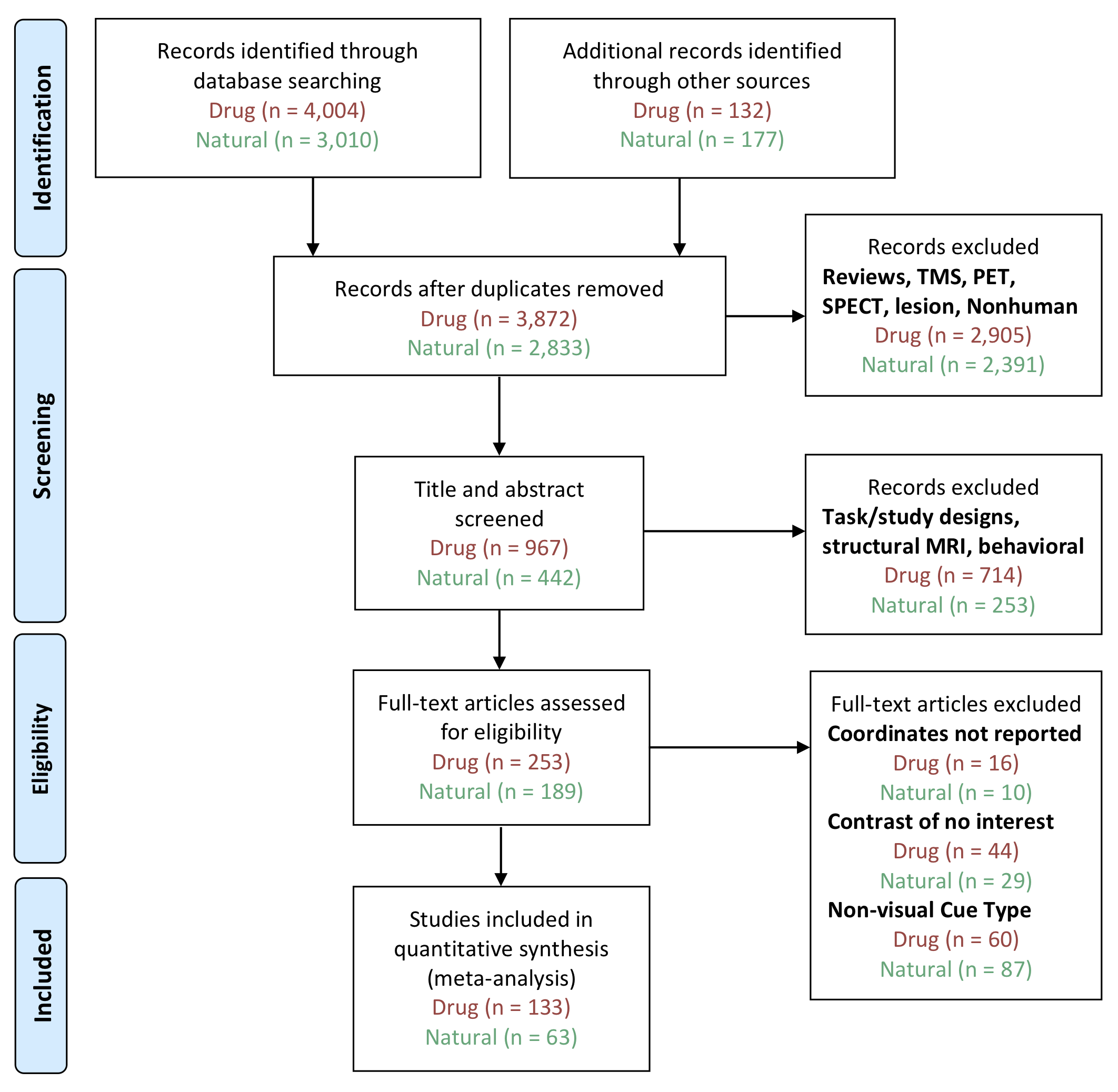

**Figure S1.** **PRISMA flow diagram.** PRISMA flow diagram of literature search and study inclusion. The chart includes counts for both drug cue-reactivity (red) and natural cue-reactivity (green) studies.

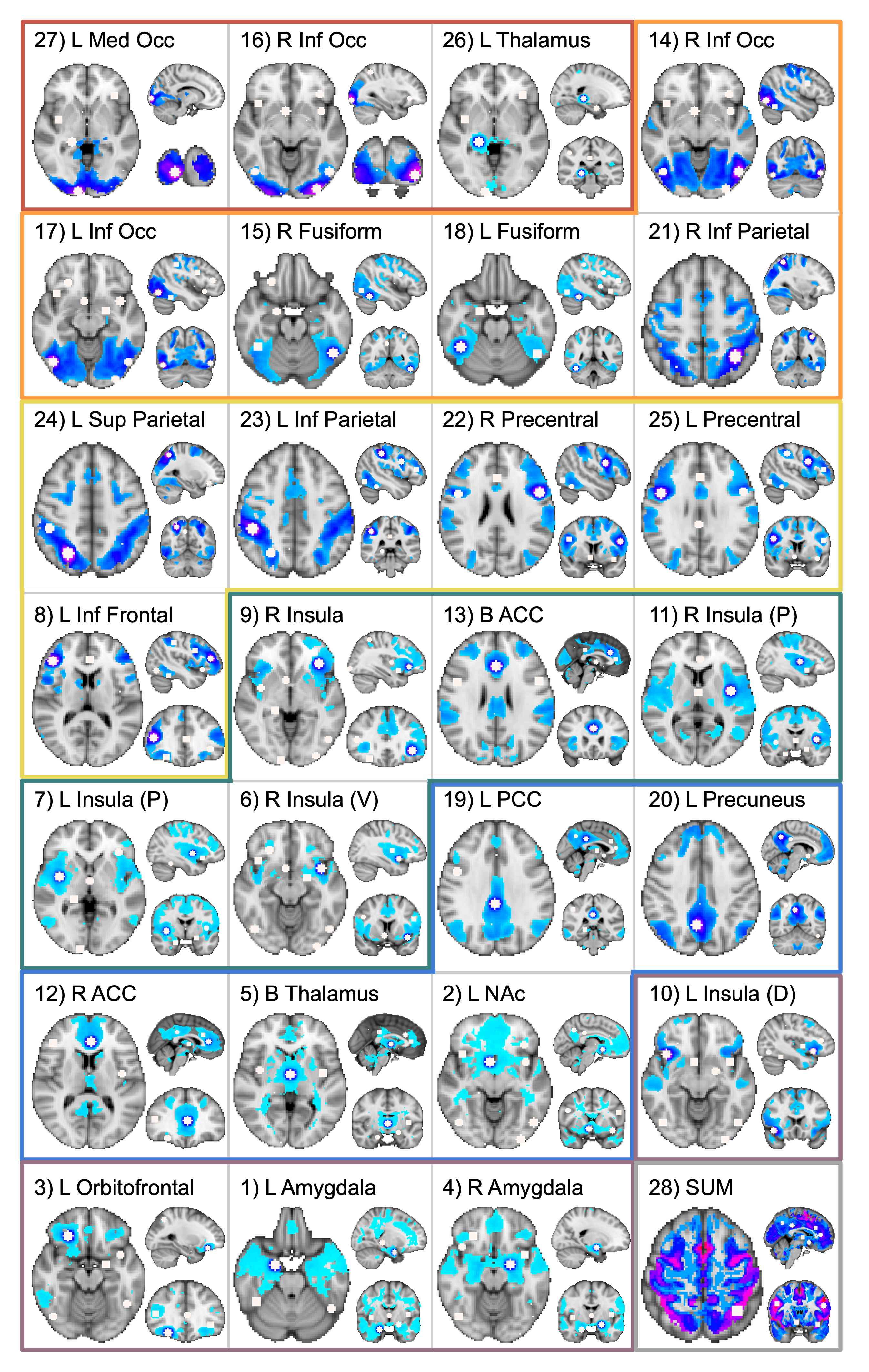

**Figure S2.** **rsFC maps for identified cue-related regions.** Resting-state functional connectivity maps for all ROIs from the *pooled* cue-reactivity meta-analytic results. Border colors represent associated cliques from *rsFC only* hierarchical clustering analysis. In relation to prior work, Clique 1 (red) containing primary visual regions, has been shown to be a consistent marker differentiating drug and neutral cues in substance users, highlighting the potential importance of patterns of activation within visual regions [1]. Clique 2 (orange) comprised of regions involving visual association, is supported by previous work demonstrating higher visual cortices are influenced by value-based learning history of the stimuli, where user’s response to drug-related stimuli is stronger than control stimuli because of reward history [2, 3]. Clique 3 (yellow) contains regions involved in cognitive control and executive functions that is supported through previous work associating impairments in cognitive control to treatment adherence across a range of tasks and substances [4]. The anterior cingulate and insula, regions involved in salience detection, compose Clique 4 (green) and have been explained to be involved in the embodiment of “feelings” from the body, as well as the initiation of behaviors across cognitive, behavioral, and affective contexts [5]. Clique 5 (blue) contains default mode network regions (PCC and ventral ACC), as well as regions involved in reward (NAc and thalamus), supporting a role in internal aspects of reward like valuation [6]. It is commonly accepted that the amygdala features a prominent role in emotion processing [7], and its connections with the orbitofrontal cortex plays a critical role in recognition and perception of emotion response, and fear extinction (Clique 6, purple) [8, 9].

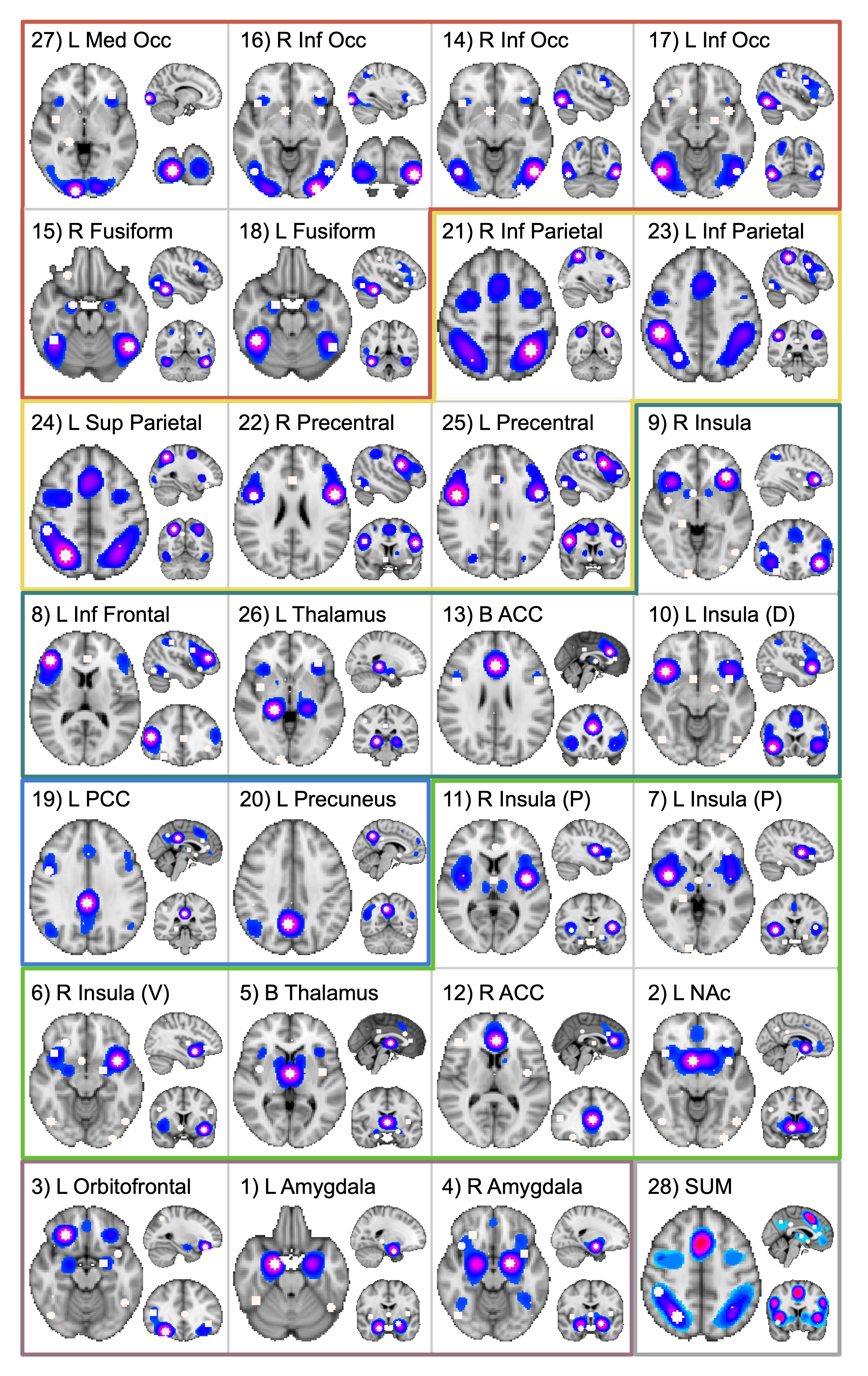

**Figure S3.** **MACM maps for identified cue-related regions.** Meta-analytic co-activation maps for all ROIs from the *pooled* cue-reactivity meta-analytic results. Border colors represent associated cliques from *MACM only* hierarchical clustering analysis. Largely similar to the rsFC maps, Clique 1 (red) is comprised of visual processing regions shown to be important for initial stages of cue stimulus differentiation [1], Clique 2 (yellow) has been shown to be involved in executive functioning associated with cognitive control mechanisms over various task paradigms and disorders of addiction [4], and Clique 6 (purple) engaged in the recognition and perception of emotion [8]. The final three cliques (green, lime green, and blue) shuffle slightly from rsFC, but provide a similar interpretation of each. rsFC Clique 4 (green) divides into two distinct cliques in the MACM hierarchical clustering analysis, where more anterior insula and frontal regions comprised Clique 3 (green), and more posterior insula and limbic regions comprised Clique 5 (lime green), both involved in salience attribution. Clique 3 (green) has been shown to be involved in the maintenance of a sense of self and attention, and Clique 5 (lime green) has been shown to be involved in the integration of salient emotional and environmental stimuli from higher-order sensory regions [5]. Finally, Clique 4 (blue) contains a primary node of the default mode network, the PCC, important for internally directed cognition and regulating the focus of attention [10].

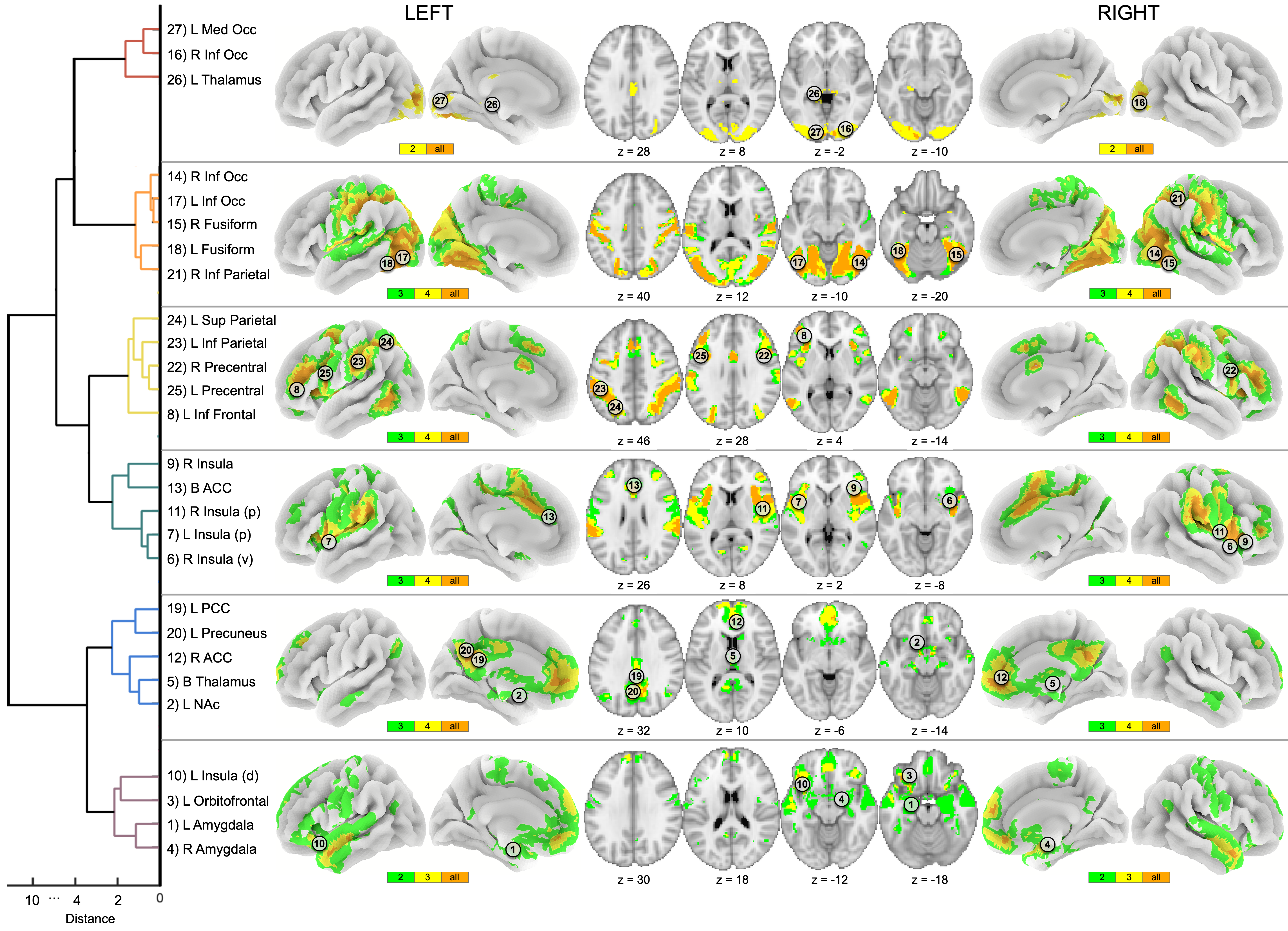

**Figure S4.** **rsFC hierarchical clustering solution.** Hierarchical clustering of the rsFC profiles of each ROI provided a clustering solution comprised of six cliques, with connectivity profiles starting from Cliques 1 and 2 originating in occipital regions and shifting to more parietal-frontal regions in Clique 3, and finally shifting to more medial-limbic regions in Cliques 4, 5, and 6. Clique 1 (red) included the L medial occipital, R inferior occipital, and L thalamus where all regions’ rsFC profiles demonstrated overlap in bilateral regions of the occipital lobe. Clique 2 (orange) included the R and L inferior occipital, R and L fusiform, and R inferior parietal where all regions’ rsFC profiles demonstrated overlap primarily in regions of occipital lobe, and precentral and postcentral gyri. Clique 3 (yellow) included the L superior and inferior parietal, R and L precentral, and L inferior frontal where all regions’ profiles demonstrated overlap in parietal, middle frontal gyrus, and cingulate regions. Clique 4 (green) included multiple regions of the R and L insula, and bilateral ACC where all regions’ rsFC profiles demonstrated overlap in the bilateral insula and cingulate gyrus. Clique 5 (blue) included the L PCC/precuneus, R ACC, bilateral thalamus, and L NAc where all regions’ rsFC profiles demonstrated overlap in medial frontal and limbic regions including the thalamus. Finally, Clique 6 (purple) included the L insula, L orbitofrontal, and R and L amygdala where all regions’ rsFC profiles demonstrated overlap in temporal and ventromedial prefrontal regions. Orange clusters on the brains demonstrate the highest degree of overlap (all regions), yellow the second degree of overlap, and green the lowest degree of overlap between rsFC maps within a clique. The horizontal axis represents the dissimilarity (or variance) between cliques, where distance is represented by Ward’s linkage algorithm.

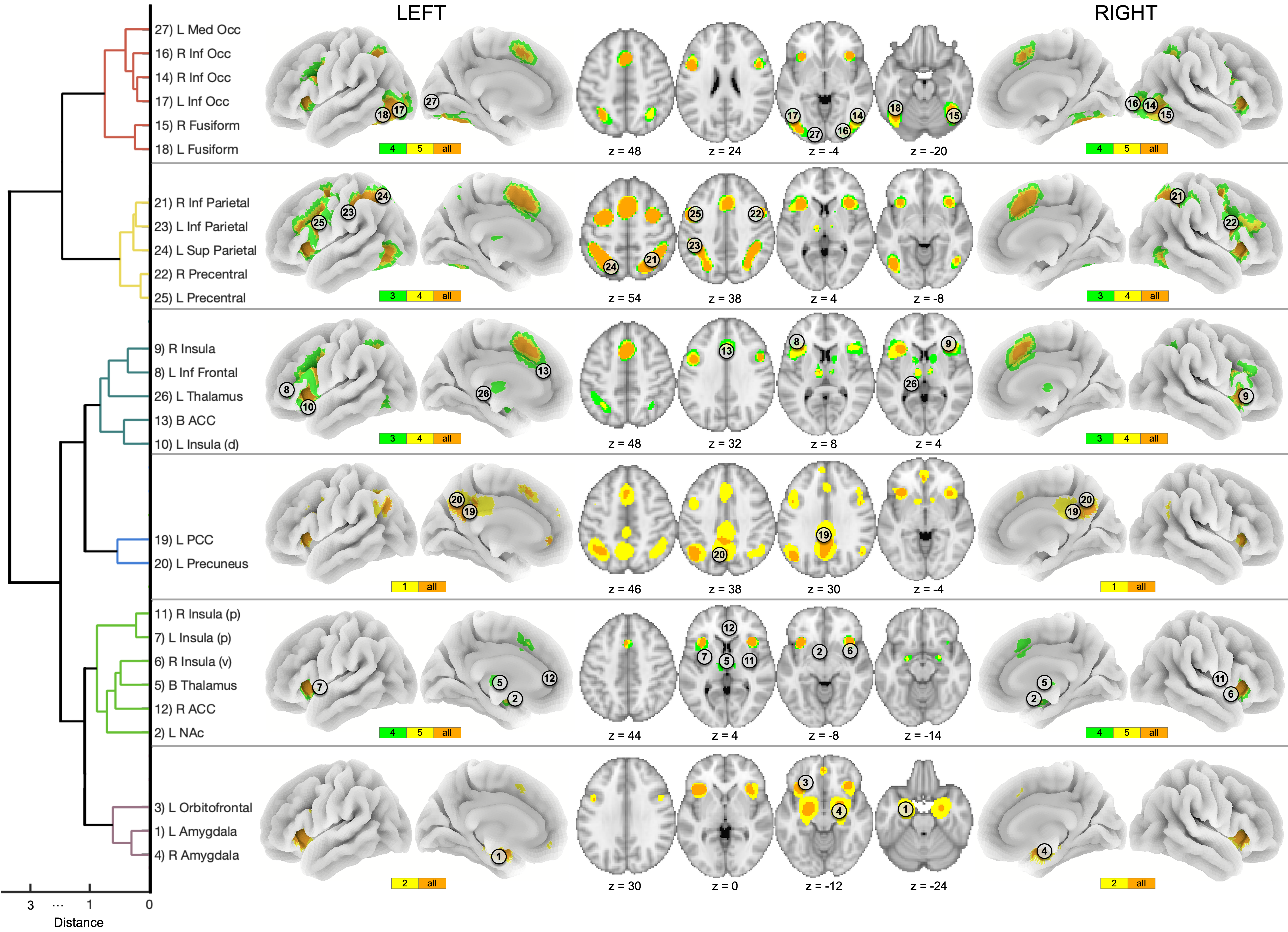

**Figure S5.** **MACM hierarchical clustering solution.** Hierarchical clustering of the MACM maps of each ROI provided a similar clustering solution of six cliques as the rsFC profiles. Similar, yet more sparse, to the rsFC results, the MACM clustering solution revealed connectivity profiles originating in occipital regions in Clique 1 to more parietal-frontal regions in Clique 2, and to more medial-limbic regions in Cliques 3 through 6. Although similar to rsFC, a few notable distinctions include: the inclusion of the R and L inferior occipital, and R and L fusiform into Clique 1 (red) rather than forming their own cluster in rsFC Clique 2 (orange), R inferior parietal clustered with other parietal/precentral regions in Clique 2 (yellow) rather than rsFC Clique 2 (orange), rsFC Clique 4 (green) split into two clusters Clique 3 (green) and Clique 5 (lime green), and PCC and precuneus clustered on their own to form Clique 5 (blue). Orange clusters on the brains demonstrate the highest degree of overlap (all regions), yellow the second degree of overlap, and green the lowest degree of overlap between MACM maps within a clique. The horizontal axis represents the dissimilarity (or variance) between cliques, where distance is represented by Ward’s linkage algorithm.

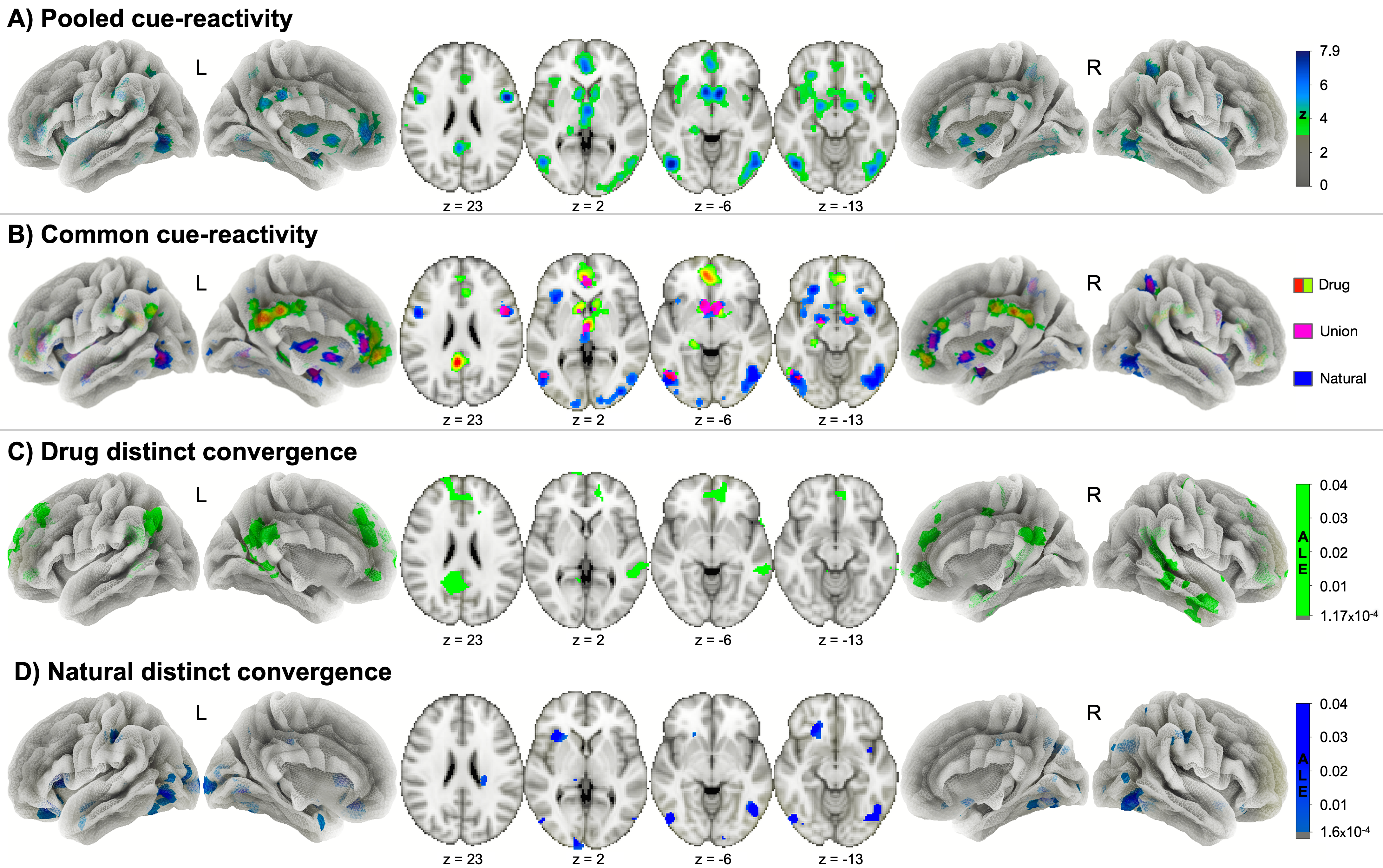

**Figure S6.** **Cue-related ALE meta-analysis results using only coordinates from whole-brain assessments.** To rule out the possibility that the inclusion of small volume corrected (SVC) coordinates biased the meta-analytic outcomes in the main text (Fig. 1), we conducted the same analyses with SVC foci removed (i.e., whole brain articles only) (*p*_cluster-corrected_ <0.05, *p*_voxel_ <0.001). The outcomes and conclusions are largely the same, and therefore all coordinates were included to be as inclusive and possible.
