## Supplemental Tables for "The cue-reactivity paradigm: An ensemble of networks driving attention and cognition when viewing drug and natural reward-related stimuli"

**SUPPLEMENTAL CONTENT**

SUPPLEMENTAL TABLES

**Table S1.** Metadata of drug cue-reactivity articles

**Table S2.** Metadata of natural cue-reactivity articles

**Table S3.** Neurosynth functional decoding

SUPPLEMENTAL REFERENCES

**Table S1.** Metadata of **drug** cue-reactivity articles included in the meta-analytic assessments.

| **No.** | **Reference** | **User n (male/female)** | **User Age (SD)** | **Cue Type** | **Imaging Modality** | **Processing** | **# Foci** | **Contrasts** | **# Contrasts** | **Analytic Procedure** |
| --- | --- | --- | --- | --- | --- | --- | --- | --- | --- | --- |
| 1 | Brumback et al. (2015) | 22 (10M/12F) | 17.93 (.71) | Alcohol | 3T fMRI | AFNI | 6 | heavy drinkers vs. control for alcohol vs. non-alcohol | 1 | WB/SVC |
| 2 | Sjoerds et al. (2014) | 30 (16M/14F) | 46.5 (8.5) | Alcohol | 3T fMRI | SPM8 | 25 | alcohol dependents: alcohol vs. neutral pictures | 1 | WB |
| 3 | Dager et al. (2014) | 43 (20M/23F) | 18.48 | Alcohol | 3T fMRI | SPM5 | 5 | moderate vs. heavy drinkers: alcohol vs. non-alcohol images | 1 | WB |
| 4 | Lukas et al. (2013) | 15 (11M/4F) | 49.93 (6.22) | Alcohol | 3T fMRI | FSL | 6 | alcohol vs. non-alcohol pictures | 1 | WB |
| 5 | Schact et al. (2013) | 48 (37M/11F) | 48.2 | Alcohol | 3T fMRI | SPM8 | 10 | alcoholic vs. nonalcoholic images | 1 | WB |
| 6 | Dager et al. (2013) | 35 (18M/17F) | 19.25 | Alcohol | 3T fMRI | SPM5 | 12 | heavy and light drinkers: alcohol vs. non-alcohol images | 2 | WB |
| 7 | Vollstadt-Klein et al. (2012) | 38 | 46 (9) | Alcohol | 3T fMRI | SPM5 | 22 | alcohol vs. neutral cues | 1 | WB |
| 8 | Claus et al. (2011) | 326 (228M/98F) | 32.5 | Alcohol | 3T fMRI | FSL | 17 | alcohol vs. control cues | 2 | WB/SVC |
| 9 | Ihssen et al. (2011) | 11 (10M/1F) | 26.91 (6.28) | Alcohol | 3T fMRI | Brain Voyager | 3 | alcohol vs. neutral | 1 | WB |
| 10 | Vollstadt-Klein et al. (2011) | 15 (9M/6F) | 50 (8) | Alcohol | 3T fMRI | SPM5 | 13 | alcohol vs. neutral cues | 1 | WB |
| 11 | Myrick et al. (2010) | 30 (22M/8F) | 28.85 | Alcohol | 3T fMRI | SPM5 | 15 | alcohol vs. beverage | 2 | WB |
| 12 | Ray et al. (2010) | 10 (5M/5F) | 20 | Alcohol | 3T fMRI | FSL | 9 | alcohol cues vs. neutral | 1 | WB |
| 13 | Vollstadt-Klein et al. (2010) | 31 (19M/12F) | 46.5 | Alcohol | 3T fMRI | SPM5 | 42 | favorite drinks vs. neutral cues | 2 | WB/SVC |
| 14 | Myrick et al. (2008) | 90 (64M/26F) | 25.81 | Alcohol | 3T fMRI | SPM2 | 24 | placebo: alcohol image vs. beverage or visual control | 2 | WB |
| 15 | Heinz et al. (2007) | 12 (6M/6F) | 40 (8) | Alcohol | 1.5T fMRI | SPM5 | 6 | alcoholics vs. controls: alcohol related vs. neutral pictures | 1 | WB |
| 16 | Park et al. (2007) | 9 (8M/1F) | 23.22 (2.48) | Alcohol | 3T fMRI | SPM99 | 11 | AUD vs. control: alcohol vs. beverage cue | 2 | WB |
| 17 | Wrase et al. (2007) | 16 (16M) | 42.4 (7.5) | Alcohol | 1.5T fMRI | SPM2 | 10 | alcoholics vs. controls: alcohol vs. neutral pictures | 2 | WB |
| 18 | Hermann et al. (2006) | 10 (10M) | 40 (7) | Alcohol | 1.5T fMRI | SPM99 | 10 | alcoholics vs. controls: alcohol associated vs. control pictures | 1 | WB |
| 19 | Myrick et al. (2004) | 10 (8M/2F) | 33.6 (11.51) | Alcohol | 1.5T fMRI | SPM96 | 15 | alcoholics: alcohol vs. beverage or control | 2 | WB |
| 20 | Tapert et al. (2004) | 8 (8F) | 19.5 (1.13) | Alcohol | 1.5T fMRI | AFNI | 12 | alcohol dependent vs. light social: alcohol vs. neutral cues | 1 | WB |
| 21 | Tapert et al. (2003) | 15 (9M/6F) | 16.96 (.78) | Alcohol | 1.5T fMRI | AFNI | 16 | AUD vs. control: alcohol beverage vs. neutral pictures | 1 | WB |
| 22 | Wrase et al. (2002) | 44 (24M/20F) | 43.5 (8.7) | Alcohol | 1.5T fMRI | SPM99 | 16 | alcohol associated vs. neutral cues | 1 | WB |
| 23 | Braus et al. (2001) | 4 (2M/2F) | 39 (6) | Alcohol | 1.5T fMRI | Brain Voyager | 3 | alcohol vs. control stimuli | 1 | WB |
| 24 | George et al. (2001) | 10 (8M/2F) | 29.9 (9.9) | Alcohol | 1.5T fMRI | SPM99 | 5 | alcohol vs. beverage or rest | 2 | WB |
| 25 | Nguyen-Louie et al. (2018) | 51 (29M/22F) | 13.7 (0.7) | Alcohol | 3T fMRI | AFNI | 1 | alcohol cues vs. neutral cues | 1 | WB |
| 26 | Kim et al. (2014) | 38 (27M/11F) | 41.6 (7.1) | Alcohol | 1.5T fMRI | Brain Voyager | 1 | patients vs. healthy: craving cues or aversion cues | 1 | WB/SVC |
| 27 | Jorde et al. (2014) | 81 (57M/24F) | 45.59 (10.33) | Alcohol | 3T fMRI | SPM5 & SPM8 | 3 | alcohol vs. neutral cues | 1 | WB/SVC |
| 28 | Fryer et al. (2013) | 44 (34M/10F) | 41.77 | Alcohol | 3T fMRI | AFNI & SPM5 | 2 | alcohol distractor vs. non-alcoholic distractor associated w/craving ratings | 1 | WB/SVC |
| 29 | Beck et al. (2012) | 46 (30M/16F) | 40.37 (6.68) | Alcohol | 1.5T fMRI | SPM8 | 7 | abstainers vs. controls vs. relapsers: alcohol associated cues vs. neutral cues | 1 | SVC |
| 30 | Schacht et al. (2011) | 9 (6M/4F) | 34.7 (12.4) | Alcohol | 3T fMRI | SPM5 | 4 | alcohol vs. neutral cues | 1 | SVC |
| 31 | Grusser et al. (2004) | 10 (5M/5F) | 36 (11) | Alcohol | 1.5T fMRI | SPM99 | 18 | alcohol-dependent vs. control: alcohol cue vs. neutral cue | 1 | SVC |
| 32 | Heinz et al. (2004) | 11 (11M) | 44.5 (6.5) | Alcohol | 1.5T fMRI | SPM99 | 4 | alcohol vs control stimuli associated w/D2 receptor availability | 1 | SVC |
| 33* | Hanlon et al. (2018) | 53 (39M/14F) | 29.9 (9.7) | Alcohol | 3T fMRI | SPM12 | 4 | all groups or alcohol: drug/alcohol vs. neutral cue | 2 | SVC |
| 34 | Beck et al. (2018) | 23 (16M/17F) | 46.17 (6.15) | Alcohol | 3T fMRI | SPM12 | 1 | alcohol dependent vs. control: alcohol-related vs. neutral cue | 1 | WB |
| 35 | Logge et al. (2019) | 30 (20M/10F) | 49.34 | Alcohol | 3T fMRI | SPM12 | 2 | alcohol vs. control images | 1 | WB |
| 36 | Bach et al. (2020) | 50 (50M) | 45.6 (8.9) | Alcohol | 3T fMRI | SPM12 | 5 | patients vs. controls: alcohol vs. neutral cues | 1 | WB |
| 37 | Fukushima et al. (2020) | 24 (17M/7F) | 47.5 (8.6) | Alcohol | 1.5T fMRI | SPM12 | 9 | alcohol use disorder vs. control: alcohol vs. non-alcohol images | 2 | WB |
| 38 | Huang et al. (2018) | 11 (8M/3F) | 48.12 (8.54) | Alcohol | 3T fMRI | SPM8 | 15 | alcohol vs. non-alcohol cues | 1 | WB |
| 39 | Koopman et al. (2019) | 41 (30M/11F) | 45.4 (10.4) | Alcohol | 3T fMRI | SPM5 | 3 | alcohol vs. neutral cues | 1 | WB |
| 1 | Wetherill et al. (2014) | 20 (12M/8F) | 29.1 (9.7) | Cannabis | 3T fMRI | SPM8 | 3 | activations to backward-masked task: cannabis vs. neutral cues | 1 | WB |
| 2 | Ames et al. (2013) | 13 (11M/2F) | 21.15 (1.9) | Cannabis | 3T fMRI | SPM8 | 5 | group by condition interaction: marijuana vs. control | 1 | WB |
| 3 | Charboneau et al. (2013) | 16 (5M/11F) | 23.7 (3.9) | Cannabis | 3T fMRI | SPM5 | 25 | cannabis cues vs. baseline or nature or food | 3 | WB |
| 4 | Fibley et al. (2009) | 38 (31M/7F) | 23.74 (7.25) | Cannabis | 3T fMRI | FSL | 46 | marijuana cue vs. control cue | 1 | WB |
| 5 | Filbey et al. (2016) | 53 (33M/20F) | 30.66 (7.48) | Cannabis | 3T fMRI | SPM8 | 24 | cannabis cues vs. neutral cues | 1 | WB |
| 6 | Machielsen et al. (2018) | 30 (30M) | 22.3 | Cannabis | 3T fMRI | SPM8 | 3 | cannabis cues vs. neutral cues | 1 | SVC |
| 7 | Goldman et al. (2013) | 12 (10M/2F) | 37.6 (10.7) | Cannabis | 3T fMRI | SPM2 | 4 | marijuana cues vs. non-marijuana cues | 1 | SVC |
| 8 | Cousijn et al. (2013) | 31 (20M/11F) | 21.3 (2.3) | Cannabis | 3T fMRI | FSL | 10 | frequent users, sporadic users, controls: cannabis vs. neutral cues | 1 | SVC |
| 9 | Zhou et al. (2019) | 26 (26M) | 22.94 (2.71) | Cannabis | 3T fMRI | SPM12 | 25 | dependent users vs. control: cannabis vs. neutral cue | 1 | WB |
| 10 | Karoly et al. (2019) | 40 (21M/19F) | 18.83 (0.96) | Cannabis | 3T fMRI | FSL | 19 | cannabis vs. non-cannabis cue | 1 | WB |
| 1 | Prisciandaro et al. (2014) | 38 (33M/5F) | 45.91 | Cocaine | 3T fMRI | SPM8 | 30 | cocaine vs. neutral cues | 2 | WB |
| 2 | Prisciandaro et al. (2013) | 30 (25M/5F) | 44.8 | Cocaine | 3T fMRI | SPM8 | 6 | cocaine vs. neutral cues | 1 | WB |
| 3 | Prisciandaro et al. (2013) | 25 (23M/2F) | 46.25 | Cocaine | 3T fMRI | SPM8 | 39 | cocaine vs. neutral cues | 2 | WB |
| 4 | Goldstein et al. (2010) | 13 (12M/1F) | 46.2 (8.7) | Cocaine | 4T fMRI | SPM2 | 6 | drug word task: control vs. CUD | 1 | WB |
| 5 | Goldstein et al. (2009) | 15 (12M/3F) | 43.6 (1.4) | Cocaine | 4T fMRI | SPM2 | 2 | cocaine addicted: drug vs. neutral words | 1 | WB |
| 6 | Garavan et al. (2000) | 17 (14M/3F) | 34 | Cocaine | 1.5T fMRI | AFNI | 19 | cocaine user exposure to film scenes of cocaine use | 1 | WB |
| 7 | Young et al. (2014) | 20 (20M) | 41.1 | Cocaine | 3T fMRI | SPM8 | 17 | baclofen treated vs. placebo: response to cocaine cues | 1 | WB/SVC |
| 8 | Goudriaan et al. (2013) | 13 | 37.7 (9.5) | Cocaine | 3T fMRI | SPM8 | 14 | cocaine vs. neutral pictures under modafinil or placebo | 2 | SVC |
| 9 | Wilcox et al. (2011) | 14 | 37.14 (8.97) | Cocaine | 3T fMRI | AFNI | 8 | cocaine users vs. healthy controls: cocaine vs. food video | 1 | WB/SVC |
| 10 | Schulte et al. (2019) | 38 (38M) | 38.59 | Cocaine | 3T fMRI | SPM12 | 15 | cocaine vs. neutral cues | 1 | WB |
| 11 | Kaag et al. (2018) | 59 (59M) | 31.4 (7.6) | Cocaine | 3T fMRI | SPM8 | 16 | cocaine vs. neutral cues | 1 | WB |
| 12 | Mayer et al. (2020) | 37 (24M/13F) | 38.15 | Cocaine | 3T fMRI | AFNI | 12 | cocaine vs. food videos | 1 | WB |
| 13 | Denomme & Shane (2020) | 20 | 37.5 (8.92) | Cocaine | 3T fMRI | SPM12 | 25 | cocaine dependent vs. control: drug vs. food-related cues | 2 | WB |
| 14 | Zhang et al. (2020) | 23 (17M/6F) | 42.4 (7.6) | Cocaine | 3T fMRI | SPM8 | 7 | cocaine vs. neutral cues | 1 | WB |
| 15* | MacNiven et al. (2018) | 36 (34M/2F) | 43.4 (13.3) | Cocaine | 3T fMRI | AFNI | 14 | drug vs. neutral cues | 2 | WB/SVC |
| 16* | Hanlon et al. (2018) | 55 (38M/17F) | 42.7 (9.6) | Cocaine | 3T fMRI | SPM12 | 3 | cocaine vs. neutral cues | 1 | SVC |
| 1 | Walter et al. (2015) | 27 (19M/8F) | 41.1 (6.1) | Heroin | 3T fMRI | SPM8 | 10 | patients vs. controls: drug-related vs. neutral cues | 2 | WB/SVC |
| 2 | Li et al. (2013) | 14 (14M) | 35.0 (6.4) | Heroin | 3T fMRI | SPM5 | 14 | heroin vs. healthy: heroin related cues vs. neutral cues | 1 | WB |
| 3 | Li et al. (2013) | 37 | 33.4 | Heroin | 3T fMRI | SPM8 | 26 | long vs. short abstinence: heroin vs. neutral cue | 2 | WB |
| 4 | Li et al. (2012) | 24 (24M) | 32.8 (6.6) | Heroin | 3T fMRI | SPM8 | 81 | heroin dependent: heroin-related vs. neutral cues | 2 | WB |
| 5 | Lou et al. (2012) | 34 (34M) | 32.4 | Heroin | 1.5T fMRI | SPM5 | 15 | heroin related cues vs. neutral cues | 1 | WB |
| 6 | Liu et al. (2011) | 16 (16M) | 36.7 (7.1) | Heroin | 3T fMRI | SPM5 | 17 | non-drug users vs. heroin users: heroin cues vs. neutral | 1 | WB |
| 7 | Mei et al. (2010) | 15 (14M/1F) | 33.5 (7.9) | Heroin | 1.5T fMRI | SPM2 | 28 | early vs. late scans: heroin-related vs. neutral cues | 1 | WB |
| 8 | Yang et al. (2009) | 15 (13M/2F) | 32.2 (3.8) | Heroin | 1.5T fMRI | SPM2 | 27 | interaction between cue-type factor and subject type factor | 1 | WB |
| 9 | Zijlstra et al. (2009) | 12 (12M) | 42.8 (5.2) | Heroin | 3T fMRI | SPM-EPI | 22 | heroin stimuli vs. baseline | 1 | WB |
| 10 | Xiao et al. (2006) | 14 (14M) | 33.2 | Heroin | 1.5T fMRI | SPM2 | 8 | drug cues vs. neutral | 1 | WB |
| 11 | Wei et al. (2020) | 61 (61M) | 36.4 | Heroin | 3T fMRI | SPM8 | 21 | heroin-related vs. neutral cues | 1 | WB |
| 12 | Zeng et al. (2018) | 37 (24M/13F) | 41.79 (2.36) | Heroin | 3T fMRI | SPM12 | 2 | drug-related vs. neutral cues | 1 | WB |
| 13 | Li et al. (2015) | 44 (44M) | 35.2 | Heroin | 3T fMRI | SPM8 | 32 | heroin-dependent vs. control: heroin-related vs. neutral cue | 1 | WB |
| 1 | Cortese et al. (2015) | 17 (13M/4F) | 29.6 (9.4) | Nicotine | 3T fMRI | FSL | 6 | no odor: smoking vs. neutral cues | 1 | WB |
| 2 | Falcone et al. (2016) | 69 (38M/31F) | 42.5 (13) | Nicotine | 3T fMRI | FSL | 6 | smoking cue minus neutral image contrast | 1 | WB |
| 3 | Claus et al. (2013) | 132 (80M/52F) | 31.36 | Nicotine | 3T fMRI | FSL | 11 | smoking vs. food videos | 1 | WB |
| 4 | Hayashi et al. (2013) | 10 (7M/3F) | 23 (3) | Nicotine | 1.5T fMRI | FSL | 15 | smoking minus control cue | 1 | WB |
| 5 | Kushnir et al. (2013) | 18 (12M/6F) | 31.1 (9.7) | Nicotine | 3T fMRI | SPM5 | 9 | pre and post-smoking: smoking vs. neutral cue | 2 | WB |
| 6 | Yalachkov et al. (2013) | 15 (7M/8F) | 28.3 (3.7) | Nicotine | 3T fMRI | Brain Voyager | 3 | smoking-related objects versus control objects | 1 | WB |
| 7 | Westbrook et al. (2013) | 47 (32M/15F) | 45 (11.35) | Nicotine | 3T fMRI | SPM8 | 2 | looking at smoking versus neutral images | 1 | WB |
| 8 | Xu et al. (2012) | 18 (18M) | 25.11 (3.03) | Nicotine | 3T fMRI | AFNI | 5 | acquaintance: cigarette vs. pen | 1 | WB |
| 9 | Kang et al. (2012) | 25 (25M) | 25 (1.2) | Nicotine | 3T fMRI | SPM5 | 17 | smoking vs. neutral cues | 1 | WB |
| 10 | Janes et al. (2012) | 24 (24F) | 44.6 | Nicotine | 3T fMRI | Brain Voyager | 11 | smoking vs. neutral images | 1 | WB |
| 11 | Franklin et al. (2011) | 22 (16M/6F) | 36.1 (2.2) | Nicotine | 3T fMRI | SPM5 | 8 | varenicline or placebo at time 1, time2: smoking vs. nonsmoking cue | 1 | WB |
| 12 | Franklin et al. (2011) | 26 (19M/7F) | 37.6 (2.3) | Nicotine | 3T fMRI | SPM | 9 | number of repeat carriers: smoking cue vs. nonsmoking cue | 1 | WB |
| 13 | Hartwell et al. (2011) | 32 (14M/18F) | 33.5 (11.5) | Nicotine | 3T fMRI | FSL | 18 | crave or resist condition: smoking vs. neutral cue | 1 | WB |
| 14 | Vollstadt-Klein et al. (2011) | 22 (22M) | 31 (7) | Nicotine | 1.5T fMRI | SPM5 | 13 | tobacco vs. control ads | 1 | WB |
| 15 | Wagner et al. (2011) | 17 (5M/12F) | 23.1 | Nicotine | 3T fMRI | SPM8 | 17 | smoking vs. nonsmoking scenes | 1 | WB |
| 16 | Wilson et al. (2012) | 90 (51M/39F) | 29.9 (7.5) | Nicotine | 3T fMRI | AFNI | 12 | cigarette vs. control cues | 1 | WB |
| 17 | Zhang et al. (2011) | 22 (11M/11F) | 31 (9.4) | Nicotine | 3T fMRI | AFNI | 6 | smoker vs. control: smoking vs. neutral cues | 1 | WB |
| 18 | Goudriaan et al. (2010) | 18 (18M) | 33.8 (9.1) | Nicotine | 3T fMRI | SPM2 | 29 | FTND-high vs. control: smoking-related vs. neutral or baseline | 2 | WB |
| 19 | Janes et al. (2010) | 21 (21F) | 46.05 | Nicotine | 3T fMRI | Brain Voyager | 48 | slip vs. abstinence: smoking vs. neutral cues | 1 | WB |
| 20 | Janes et al. (2010) | 28 (28F) | 44.3 (10.2) | Nicotine | 3T fMRI | Brain Voyager | 4 | smoking vs. neutral images | 1 | WB |
| 21 | King et al. (2010) | 12 (10M/2F) | 23.2 (1.8) | Nicotine | 3T fMRI | SPM2 | 4 | smoking content vs. control content | 1 | WB |
| 22 | Janes et al. (2009) | 13 (13F) | 43.2 (11.5) | Nicotine | 3T fMRI | Brain Voyager | 70 | no abstinence or abstinence: smoking related vs. neutral images | 2 | WB |
| 23 | Dagher et al. (2009) | 15 (8M/7F) | 29 | Nicotine | 3T fMRI | fmri-stat | 11 | stress vs. nonstress: smoking vs. control video | 1 | WB |
| 24 | Artiges et al. (2009) | 13 (8M/5F) | 26 (4) | Nicotine | 1.5T fMRI | SPM2 | 2 | smokers vs. nonsmokers: smoking vs. neutral cues | 1 | WB |
| 25 | McClernon et al. (2009) | 18 (7M/11F) | 28.6 (7.5) | Nicotine | 4T fMRI | SPM5 | 19 | abstinent: smoking vs. control cues | 1 | WB |
| 26 | Brody et al. (2007) | 42 (30M/12F) | 38 (12.4) | Nicotine | 1.5T fMRI | FSL | 41 | crave vs. resist: cigarette vs. neutral cues | 2 | WB |
| 27 | McBride et al. (2006) | 20 (10M/10F) | 27 (8) | Nicotine | 1.5T fMRI | fmri-stat | 10 | smoking related vs. neutral cues | 1 | WB/SVC |
| 28 | Smolka et al. (2006) | 10 (10M) | 32 (5) | Nicotine | 1.5T fMRI | SPM99 | 14 | smoking cues vs. neutral cues assoc. w/ severity | 1 | WB |
| 29 | David et al. (2005) | 9 (4M/5F) | 34.4 | Nicotine | 3T fMRI | FSL | 3 | smoking vs. neutral cues | 1 | WB |
| 30 | Lee et al. (2005) | 8 (8M) | 17 (.76) | Nicotine | 1.5T fMRI | AFNI | 14 | 2D or 3D: smoking vs. neutral cues | 1 | WB |
| 31 | Wilson et al. (2005) | 22 (22M) | 24.4 (4.9) | Nicotine | 1.5T fMRI | AFNI | 9 | cigarette cue vs. neutral cues | 1 | WB |
| 32 | Moran et al. (2018) | 19 (9M/10F) | 31.5 (5.8) | Nicotine | 3T fMRI | FSL | 9 | control smoker vs. schizophrenia smoker: smoking vs. neutral cues | 1 | WB |
| 33 | Do & Galvan (2016) | 39 (25M/14F) | 22.29 | Nicotine | 3T fMRI | FSL | 47 | smoking cues vs. neutral cues or baseline | 2 | WB |
| 34 | Janes et al. (2015) | 17 (8M/9F) | 31.4 (6.1) | Nicotine | 3T fMRI | FSL | 9 | smoking vs. neutral cues | 1 | WB |
| 35 | Courtney et al. (2014) | 40 (25M/15F) | 31.17 (8.82) | Nicotine | 3T fMRI | FSL | 1 | cigarette cues vs. control cues | 1 | SVC |
| 36 | Ko et al. (2013) | 16 (16M) | 25.38 (3.36) | Nicotine | 3T fMRI | SPM5 | 15 | smoking vs. neutral cues | 1 | WB/SVC |
| 37 | Rubinstein et al. (2011) | 12 (7M/5F) | 16.3 (.98) | Nicotine | 3T fMRI | MATLAB | 15 | smoker vs. nonsmoker: smoking cue vs. neutral cues | 2 | WB/SVC |
| 38 | David et al. (2007) | 8 (8F) | 55 (9) | Nicotine | 3T fMRI | FSL | 4 | smoking or abstinent: smoking vs. neutral cues | 1 | SVC |
| 39 | McClernon et al. (2007) | 15 (2M/13F) | 39.8 (9.94) | Nicotine | 1.5T fMRI | SPM2 | 4 | smoking vs. control cues | 1 | SVC |
| 40 | Allenby et al. (2020) | 75 (40M/35F) | 43 (12.7) | Nicotine | 3T fMRI | FSL | 6 | smoking vs. neutral cues | 1 | WB |
| 41 | Zanchi et al. (2015) | 14 (4M/10F) | 29.3 (6) | Nicotine | 3T fMRI | FSL | 2 | smokers vs. non: smoking vs. control videos | 1 | WB |
| 42 | Zhao et al. (2020) | 26 (26M) | college | Nicotine | 3T fMRI | FSL | 32 | smoking-related vs. neutral stimuli | 1 | WB |
| 43 | Bi et al. (2017) | 35 (35M) | 21.03 (1.22) | Nicotine | 3T fMRI | SPM8 | 39 | smoking vs. neutral cues | 1 | WB |
| 44 | Dinh-Williams et al. (2014) | 30 (15M/15F) | 31.9 (9.4) | Nicotine | 3T fMRI | SPM5 | 5 | appetitive cigarette vs. neutral images | 1 | WB |
| 45 | Ghahremani et al. (2018) | 21 (11M/10F) | 22.6 (2) | Nicotine | 3T fMRI | FSL | 21 | smoking vs. non-smoking cues | 2 | WB |
| 46 | Mondino et al. (2018) | 29 (9M/20F) | 41 | Nicotine | 1.5T fMRI | SPM12 | 27 | baseline: smoking vs. neutral cues | 1 | WB |
| 47 | Moran-Santa Maria et al. (2015) | 58 (35M/23F) | 34.5 (1.6) | Nicotine | 3T fMRI | FSL | 7 | smoking vs. neutral cues | 1 | WB |
| 48 | Ray et al. (2015) | 39 (24M/15F) | 31.41 (8.81) | Nicotine | 3T fMRI | FSL | 5 | cigarette vs. neutral cues | 1 | WB |
| 49 | Versace et al. (2011) | 35 (21M/14F) | 42.7 (11.3) | Nicotine | 3T fMRI | Brain Voyager | 13 | cigarette vs. neutral cues | 1 | WB |
| 50 | Wilson et al. (2013) | 82 (70M/12F) | 33 (8.3) | Nicotine | 3T fMRI | AFNI/FSL | 12 | cigarette vs. control cues | 1 | WB |
| 51 | McClernon et al. (2016) | 30 (13M/17F) | 35.4 (11.5) | Nicotine | 3T fMRI | FSL | 18 | smoking vs. non-smoking stimuli | 2 | WB/SVC |
| 52 | Janes et al. (2015) | 18 (8M/10F) | 25.06 (4.75) | Nicotine | 3T fMRI | FSL | 9 | smoking vs. neutral images | 2 | WB |
| 53 | Van Rensburg et al. (2009) | 10 (6M/4F) | 34 | Nicotine | 1.5T fMRI | SPM2 | 22 | smoking vs. neutral images | 1 | WB |
| 54 | Karch et al. (2019) | 36 (25M/11F) | 43.83 (12.37) | Nicotine | 3T fMRI | Brain Voyager | 9 | abstinent and relapse: tobacco-related vs. neutral images | 2 | WB |
| *55 | Hanlon et al. (2018) | 48 (24M/24F) | 37.1 (11.9) | Nicotine | 3T fMRI | SPM12 | 1 | cigarette vs. neutral cues | 1 | SVC |
| **133** | **TOTAL:** | **2739M/1243F** | **34.37** |  |  |  | **1870** |  | **164** |  |

**Note.** Articles organized and sub-divided by drug class. WB = whole-brain assessments, SVC = small-volume corrected assessments, M = male, F = female, FSL = FMRIB Software Library, AFNI = Analysis of Functional NeuroImages, SPM = Statistical Parametric Mapping. * = articles with experiments that fall under multiple cue type categories. One article did not report participant age, but did state college-aged, and five additional articles did not report female/male participant ratios.

**Table S2.** Metadata of **natural** cue-reactivity articles included in the meta-analytic assessments.

| **No.** | **Reference** | **n (male/female)** | **Age (SD)** | **Cue Type** | **Imaging Modality** | **Processing** | **# Foci** | **Contrasts** | **# Contrasts** | **Analytic Procedure** |
| --- | --- | --- | --- | --- | --- | --- | --- | --- | --- | --- |
| 1 | Frankfort et al. (2014) | 34 (34F) | 20 | Food | 3T fMRI | Brain Voyager | 19 | control vs. experimental: chocolate vs. neutral cue | 2 | WB |
| 2 | Kroemer et al. (2013) | 26 (13M/13F) | 24.4 (3.4) | Food | 3T fMRI | SPM5 | 16 | food pictures vs. control pictures | 1 | WB |
| 3 | Kim et al. (2012) | 58 (58F) | 23.8 | Food | 1.5T fMRI | AFNI | 35 | anorexia vs. bulimia vs. control: food vs. non-food images | 6 | WB |
| 4 | Grosshans et al. (2012) | 44 (14M/30F) | 40.7 (12.2) | Food | 3T fMRI | SPM5 | 4 | food cues vs. neutral cues | 1 | WB |
| 5 | Murdaugh et al. (2012) | 38 (11M/27F) | 46.6 | Food | 3T fMRI | SPM8 | 30 | control or obese group: high-calorie food vs. car images | 3 | WB/SVC |
| 6 | Rothemund et al. (2011) | 24 (24F) | 25 | Food | 1.5T fMRI | SPM2 | 24 | control or anorexic: high calorie, low calorie, utensil vs. neutral | 2 | WB |
| 7 | Cornier et al. (2009) | 22 (12M/10F) | 34.4 (5.1) | Food | 3T fMRI | SPM5 | 23 | thin: hedonic vs. non-food images in the eucaloric state | 1 | WB |
| 8 | Malik et al. (2008) | 20 (20M) | 23.65 | Food | 1.5T fMRI | fmri-stat | 74 | ghrelin and control: food vs. scenery stimuli | 4 | WB |
| 9 | Rothemund et al. (2007) | 26 (26F) | 30 | Food | 1.5T fMRI | SPM2 | 11 | obese or control: low calorie or high calorie vs. neutral | 3 | WB |
| 10 | Porubska et al. (2006) | 12 (7M/5F) | 27.17 (5.36) | Food | 1.5T fMRI | SPM2 | 4 | food vs. food-neutral stimuli | 1 | WB |
| 11 | Uher et al. (2006) | 18 (8M/10F) | 28.9 | Food | 1.5T fMRI | N/A | 5 | food related visual stimuli vs. nonfood stimuli | 1 | WB |
| 12 | St-Onge et al. (2005) | 12 (6M/6F) | 29.8 (1.8) | Food | 1.5T fMRI | SPM99 | 9 | food vs. nonfood stimuli | 1 | WB |
| 13 | Killgore et al. (2003) | 13 (13F) | 23.5 (2.1) | Food | 1.5T fMRI | SPM99 | 30 | low-calorie and high-calorie vs. control stimuli | 3 | WB |
| 14 | Kuhn et al. (2016) | 77 (30M/47F) | 26 (0.65) | Food | 3T fMRI | SPM8 | 17 | food vs. non-food cues | 1 | WB |
| 15 | Weygandt et al. (2012) | 67 (67F) | 24.75 | Food | 1.5T fMRI | SPM2 | 29 | eating disorder or control: food vs. neutral | 1 | SVC |
| 16 | Demos et al. (2012) | 58 (58F) | 18 | Food | 3T fMRI | SPM2 | 2 | food vs. nonfood cues | 1 | SVC |
| 17 | Scharmuller et al. (2012) | 26 (26F) | 26.1 | Food | 3T fMRI | SPM8 | 5 | obese or normal-weight: food vs neutral cues | 2 | SVC |
| 18 | Schienle et al. (2009) | 67 (67F) | 24.2 | Food | 1.5T fMRI | SPM2 | 41 | eating disorder or control: food vs. neutral | 4 | WB/SVC |
| 19 | Schur et al. (2009) | 10 (10F) | 29.4 (12.1) | Food | 3T fMRI | FSL | 23 | fattening and/or non-fattening vs. object images | 2 | SVC |
| 20 | Beaver et al. (2006) | 12 (5M/7F) | 22 (2.4) | Food | 3T fMRI | SPM99 | 16 | appetizing vs. non-food | 1 | SVC |
| 21 | Simmons et al. (2005) | 9 (3M/6F) | 31.5 | Food | 3T fMRI | SPM99 | 6 | foods vs. locations | 1 | SVC |
| 22 | Schulte et al. (2019) | 44 (44F) | 30.55 (4.04) | Food | 3T fMRI | SPM12 | 2 | food vs. household item cues | 2 | WB |
| 23 | Limbrick-Oldfield et al. (2017) | 42 (42M) | 29.5 | Food | 3T fMRI | FSL | 4 | food vs. food-matched neutral cues | 1 | WB |
| 24 | Farr & Mantzoros (2017) | 11 (6M/5F) | 45.5 (3.88) | Food | 3T fMRI | SPM8 | 2 | nondiabetic: all food vs. non-food stimuli | 1 | WB |
| 25 | King et al. (2018) | 12 (4M/8F) | 25 (6.5) | Food | 3T fMRI | SPM12 | 5 | high fat, complex carb, sugar vs. non-food images | 2 | SVC |
| 26 | Nakamura et al. (2019) | 35 (16M/19F) | 17.2 (1.9) | Food | 3T fMRI | SPM12 | 34 | food vs. non-food images | 2 | WB/SVC |
| 27 | Wiers et al. (2020) | 16 (10M/6F) | 38.81 (13.4) | Food | 3T fMRI | SPM8 | 18 | food vs. neutral cues | 4 | WB |
| 28 | Borgan et al. (2019) | 28 (22M/6F) | 26.43 (5.46) | Food | 3T fMRI | SPM12 | 9 | controls: food vs. non-food images | 2 | WB/SVC |
| 29 | Moreno-Padilla et al. (2018) | 77 (37M/40F) | 16.64 | Food | 3T fMRI | SPM8 | 21 | appetizing vs. plain food vs. baseline | 1 | WB |
| 30* | MacNiven et al. (2018) | 40 (24M/16F) | 32 (11.6) | Food | 3T fMRI | AFNI | 23 | food vs. neutral cues | 2 | WB/SVC |
| 1 | Borg et al. (2014) | 20 (20F) | 22 (2.1) | Sex | 3T fMRI | SPM8 | 23 | penetration vs. neutral objects | 1 | WB |
| 2 | Borg et al. (2014) | 62 (62F) | 23.8 | Sex | 3T fMRI | SPM8 | 31 | penetration vs. bodies | 2 | WB |
| 3 | Oei et al. (2014) | 37 (37M) | 21.96 | Sex | 3T fMRI | FSL | 20 | sexual vs. neutral or fixation | 2 | WB |
| 4 | Wetherill et al. (2014) | 20 (12M/8F) | 29.1 (9.7) | Sex | 3T fMRI | SPM8 | 5 | sexual vs. neutral cues | 1 | WB |
| 5 | Wehrum et al. (2013) | 100 (50M/50F) | 25.4 (4.8) | Sex | 1.5T fMRI | SPM8 | 26 | sex vs. neutral, positive, or negative | 2 | WB |
| 6 | Versace et al. (2013) | 27 (27F) | 54.04 (3.09) | Sex | 3T fMRI | Brain Voyager | 12 | sexual desire: emotional vs. neutral | 2 | WB |
| 7 | Kim et al. (2013) | 23 (23F) | 38.4 (10) | Sex | 3T fMRI | SPM2 | 12 | erotic vs. neutral cues | 1 | WB |
| 8 | Oei et al. (2012) | 53 (53M) | 22.7 | Sex | 3T fMRI | FSL | 48 | sexual vs. neutral or fixation | 2 | WB |
| 9 | Gillath et al. (2012) | 39 (19M/20F) | 19.65 | Sex | 1.5T fMRI | SPM5 | 28 | supraliminal or subliminal: sexual vs. neutral cues | 2 | WB |
| 10 | Kagerer et al. (2011) | 21 (21M) | 28 (4.5) | Sex | 1.5T fMRI | SPM8 | 10 | homosexual and heterosexual: sexual arousal vs. neutral | 1 | WB |
| 11 | Bianchi-Demicheli et al. (2011) | 28 (28F) | 31.1 (7.02) | Sex | 3T fMRI | SPM5 | 25 | hypoactive or no hypo desire: erotic vs. non-erotic cues | 2 | WB |
| 12 | Hu et al. (2011) | 28 (28M) | 26.5 | Sex | 1.5T fMRI | SPM2 | 11 | homosexual vs. heterosexual: heterosexual, or gay couple vs. neutral images | 2 | WB |
| 13 | Zhang et al. (2011) | 32 (32M) | 27.05 | Sex | 1.5T fMRI | SPM2 | 56 | homosexual or heterosexual: aversive sexual stim vs. rest | 2 | WB |
| 14 | Barros-Loscertales et al. (2010) | 45 (45M) | 21.82 | Sex | 1.5T fMRI | SPM2 | 19 | erotic pictures vs. neutral pictures | 1 | WB |
| 15 | Sescousse et al. (2010) | 18 (18M) | 24 (3.3) | Sex | 1.5T fMRI | SPM2 | 20 | erotic vs. neutral pictures | 1 | WB |
| 16 | Arnow et al. (2009) | 36 (36F) | 24.9 | Sex | 3T fMRI | SPM99/SPM2 | 120 | control or hypo desire: erotic vs. sports cues | 2 | WB |
| 17 | Gizewski et al. (2009) | 36 (12M/24F) | 31.33 | Sex | 1.5T fMRI | SPM99 | 14 | hetero or trans: erotic vs. neutral film excerpts | 2 | WB |
| 18 | Buhler et al. (2008) | 10 (10M) | 32 (5) | Sex | 1.5T fMRI | SPM99 | 24 | erotic vs. neutral cue | 2 | WB |
| 19 | Schiffer et al. (2008) | 12 (12M) | 36.1 (7.5) | Sex | 1.5T fMRI | SPM2 | 56 | control group: sexual vs. neutral cues | 2 | WB |
| 20 | Paul et al. (2008) | 24 (24M) | 33.4 (7.2) | Sex | 1.5T fMRI | SPM2 | 25 | homosexual or heterosexual: erotic vs. neutral stimuli | 2 | WB |
| 21 | Brunetti et al. (2008) | 18 (18M) | 24.89 | Sex | 1.5T fMRI | Brain Voyager | 26 | erotic vs. sport and neutral clips | 1 | WB |
| 22 | Safron et al. (2007) | 24 (24M) | 21 | Sex | 3T fMRI | AFNI | 15 | preferred sexual stimuli vs. sports stimuli | 1 | WB |
| 23 | Ponseti et al. (2006) | 53 (27M/26F) | 25.45 | Sex | 1.5T fMRI | SPM2 | 13 | sexual vs. nonsexual stimuli | 1 | WB |
| 24 | Moulier et al. (2006) | 10 (10M) | 21.7 | Sex | 1.5T fMRI | SPM99 | 24 | sexual stimuli vs. dressed humans | 1 | WB |
| 25 | Stark et al. (2005) | 12 (6M/6F) | 28.2 | Sex | 1.5T fMRI | SPM99 | 8 | control: erotic vs. neutral | 1 | WB/SVC |
| 26 | Ferretti et al. (2005) | 10 (10M) | 23 | Sex | 1.5T fMRI | Brain Voyager | 45 | erotic vs. sports stimuli | 2 | WB |
| 27 | Beauregard et al. (2001) | 10 (10M) | 23.5 | Sex | 1.5T fMRI | SPM99 | 18 | sexual arousal or inhibition: erotic vs. neutral cues | 2 | WB/SVC |
| 28 | Cyders et al. (2016) | 27 (14M/13F) | 25.2 (3.6) | Sex | 3T fMRI | SPM8 | 19 | sexual vs. nonsexual images | 1 | WB |
| 29 | Kuhn et al. (2014) | 64 (64M) | 28.9 (6.62) | Sex | 3T fMRI | SPM8 | 2 | sexual cue vs. fixation and nonsexual | 1 | SVC |
| 30 | Wehrum-Osinsky et al. (2014) | 56 (32M/24F) | 25.9 (5.4) | Sex | 1.5T fMRI | SPM8 | 16 | sexual vs. neutral cues | 1 | SVC |
| 31 | Kim et al. (2006) | 10 (10M) | 52 | Sex | 1.5T fMRI | SPM99 | 13 | sexual vs. neutral stimuli | 1 | SVC |
| 32 | Stark et al. (2019) | 70 (37M/33F) | 25.73 (4.62) | Sex | 3T fMRI | SPM12 | 20 | sexual vs. neutral stimuli | 2 | WB/SVC |
| 33 | Strahler et al. (2018) | 97 (47M/50F) | 25.27 | Sex | 1.5T fMRI | SPM12 | 22 | sexual vs. neutral images | 2 | SVC |
| **63** | **TOTAL:** | **972M/1138F** | **28.02** |  |  |  | **1367** |  | **110** |  |

**Note.** Articles organized and sub-divided by natural category. WB = whole-brain assessments, SVC = small-volume corrected assessments, M = male, F = female, FSL = FMRIB Software Library, AFNI = Analysis of Functional NeuroImages, SPM = Statistical Parametric Mapping.

**Table S3.** Neurosynth functional decoding outcomes for subgroups of cue-related brain regions.

| TERM TYPE | TERM RANK | **RED** | | **ORANGE** | | **YELLOW** | | **GREEN** | | **BLUE** | | **PURPLE** | |
| --- | --- | --- | --- | --- | --- | --- | --- | --- | --- | --- | --- | --- | --- |
|  |  | Term | Corr. | Term | Corr. | Term | Corr. | Term | Corr. | Term | Corr. | Term | Corr. |
| **Anatomical** | |  |  |  |  |  |  |  |  |  |  |  |  |
|  | 1 | occipito temporal (2)^1^ | 0.372 | occipito temporal (3)^4^ | 0.535 | parietal (2)^10^ | 0.534 | anterior insula (2)^14^ | 0.513 | anterior | 0.373 | amygdala (2)^27^ | 0.420 |
|  | 2 | inferior frontal | 0.332 | fusiform (5)^5^ | 0.495 | intraparietal (3)^11^ | 0.532 | insula (1)^15^ | 0.437 | anterior insula (1)^19^ | 0.352 | anterior insula (1)^28^ | 0.378 |
|  | 3 | intraparietal (2)^2^ | 0.312 | lateral occipital | 0.435 | frontal | 0.486 | inferior frontal | 0.370 | anterior cingulate (2)^20^ | 0.325 | insula | 0.332 |
|  | 4 | fusiform gyrus (1)^3^ | 0.309 | extrastriate | 0.420 | fronto parietal (1)^12^ | 0.458 | anterior | 0.344 | striatum (1)^21^ | 0.300 | amygdala hippocampus | 0.326 |
|  | 5 | inferior | 0.306 | ventral visual | 0.407 | inferior | 0.445 | frontal | 0.326 | cingulate (1)^22^ | 0.299 | anterior | 0.318 |
|  | 6 | ventral visual | 0.300 | occipital (1)^6^ | 0.387 | inferior frontal | 0.419 | frontal operculum (1)^16^ | 0.311 | mesolimbic | 0.266 | inferior frontal | 0.275 |
|  | 7 | occipital cortex | 0.295 | intraparietal (2)^7^ | 0.358 | pre sma | 0.418 | acc (1)^17^ | 0.284 | insula | 0.266 | amygdala anterior | 0.267 |
|  | 8 | frontal | 0.293 | inferior | 0.352 | posterior parietal | 0.410 | insula inferior | 0.273 | prefrontal | 0.264 | amygdala insula | 0.246 |
|  | 9 | extrastriate | 0.284 | inferior frontal | 0.349 | inferior parietal | 0.379 | inferior | 0.263 | ventral striatum (2)^23^ | 0.256 | mesolimbic | 0.235 |
|  | 10 | fronto parietal | 0.278 | inferior occipital | 0.297 | frontal eye | 0.374 | pre sma | 0.260 | caudate nucleus | 0.223 | anterior cingulate | 0.234 |
| **Functional** | |  |  |  |  |  |  |  |  |  |  |  |  |
|  | 1 | task | 0.455 | visual | 0.566 | task | 0.622 | task | 0.358 | general | 0.339 | pictures | 0.385 |
|  | 2 | visual | 0.440 | object (1)^8^ | 0.498 | demands | 0.495 | general | 0.339 | reward (1)^24^ | 0.312 | neutral | 0.383 |
|  | 3 | stimulus | 0.373 | visual word (1)^9^ | 0.473 | working memory (2)^13^ | 0.480 | stimulus (1)^18^ | 0.311 | task | 0.295 | emotional (1)^29^ | 0.346 |
|  | 4 | word | 0.371 | reading | 0.442 | attentional | 0.435 | demands | 0.295 | incentive (1)^25^ | 0.294 | stimuli | 0.341 |
|  | 5 | demands | 0.364 | orthographic | 0.437 | load | 0.408 | response | 0.287 | anticipation | 0.279 | responses | 0.329 |
|  | 6 | reading | 0.363 | characters | 0.421 | visually | 0.407 | painful | 0.275 | monetary | 0.278 | general | 0.317 |
|  | 7 | engaged | 0.354 | word form | 0.420 | target | 0.398 | conflict | 0.268 | engaged | 0.277 | unpleasant | 0.302 |
|  | 8 | orthographic | 0.350 | face | 0.420 | phonological | 0.391 | response inhibition | 0.250 | motivation (1)^26^ | 0.273 | valence | 0.301 |
|  | 9 | object | 0.339 | perceptual | 0.411 | stimulus | 0.390 | task difficulty | 0.246 | retrieval | 0.270 | aversive | 0.298 |
|  | 10 | general | 0.338 | task | 0.408 | task difficulty | 0.389 | stop | 0.242 | decision | 0.251 | affective | 0.295 |
| SUMMARY | | **VISUAL** | | **VISUAL ASSOCIATION** | | **COGNITIVE CONTROL** | | **SALIENCE** | | **VALUATION** | | **EMOTION** | |
| **(#) = number of term near-duplicates in Neurosynth output** | | | | | | | | | |  |  |  |  |
| **Superscript = corresponding near-duplicates in output** | | | | | | | | | |  |  |  |  |
| **Near-Duplicates:** | | ^1^occipito, occipitotemporal | | ^4^occipito, occipitotemporal, occipitotemporal cortex | | ^10^parietal cortex, parietal network | | ^14^insula anterior, anterior insular | | ^19^insula anterior | | ^27^cortex amygdala, amygdala responses | |
|  |  | ^2^intraparietal sulcus, sulcus ips | | ^5^fusiform gyrus, ffa, fusiform face, face ffa, fusiform gyri | | ^11^intraparietal sulcus, sulcus ips, ips | | ^15^insular cortex | | ^20^acc, cortex acc | | ^28^insula anterior | |
|  |  | ^3^fusiform | | ^6^occipital cortex | | ^12^frontoparietal | | ^16^operculum | | ^21^striatal | | ^29^emotion | |
|  |  |  |  | ^7^intraparietal sulcus, sulcus ips, | | ^13^working, memory wm | | ^17^anterior cingulate | | ^22^cingulate cortex | |  |  |
|  |  |  |  | ^8^objects | |  |  | ^18^stimuli | | ^23^nucleus accumbens, accumbens | |  |  |
|  |  |  |  | ^9^word | |  |  |  |  | ^24^rewards | |  |  |
|  |  |  |  |  |  |  |  |  |  | ^25^incentive delay | |  |  |
|  |  |  |  |  |  |  |  |  |  | ^26^motivational | |  |  |

**Note.** Neurosynth top 10 anatomical, and top 10 functional terms linked with each cue-reactivity clique. Repeats or near-duplicates of terms that appeared when identifying the unique top 10 terms are marked with a superscript, as well as a number in parenthesis with the number of repeats. A list of the corresponding near-duplicates are located in the lower portion of the table. Functional relationships of regions in Clique 1 (red) were highly associated with *visual*, *stimulus*, and *word*, and consistently linked with *occipito temporal*, *inferior frontal*, and *intraparietal* anatomical regions. The functional connections between regions of Clique 2 (orange) were consistently associated with *visual*, *object,* and *visual word*, and linked with *occipito temporal*, *fusiform*, and *lateral occipital* regions. Clique 3’s (yellow) functional profile was defined by *demands*, *working memory*, and *attentional* while anatomical regions included *parietal*, *intraparietal*, and *frontal*. Functional terms consistently associated with Clique 4 (green) included *general*, *stimulus*, and *demands*, with associated anatomical regions: *anterior insula*, *insula*, and *inferior frontal*. The functional relationships of regions in Clique 5 (blue) were highly associated with *general*, *reward*, and *incentive*, and linked with *anterior*, *anterior insula*, and *anterior cingulate* anatomical regions. Finally, regions in Clique 6 (purple) were functionally defined by *pictures*, *neutral*, and *emotional*, and linked with *amygdala*, *anterior insula*, and *insula* regions.
